# Exonic Mosaic Mutations Contribute Risk for Autism Spectrum Disorder

**DOI:** 10.1101/083428

**Authors:** Deidre R. Krupp, Rebecca A. Barnard, Yannis Duffourd, Sara A. Evans, Ryan M. Mulqueen, Raphael Bernier, Jean-Baptiste Rivière, Eric Fombonne, Brian J. O’Roak

**Author notes:** These authors contributed equally to this work (Deidre R. Krupp, Rebecca A. Barnard).

## Abstract

Genetic risk factors for autism spectrum disorder (ASD) have yet to be fully elucidated. Postzygotic mosaic mutations (PMMs) have been implicated in several neurodevelopmental disorders and overgrowth syndromes. We systematically evaluated PMMs by leveraging whole-exome sequencing data on a large family-based ASD cohort, the Simons Simplex Collection. We found evidence that 11% of published single nucleotide variant (SNV) *de novo* mutations are potentially PMMs. We then developed a robust SNV PMM calling approach that leverages complementary callers, logistic regression modeling, and additional heuristics. Using this approach, we recalled SNVs and found that 22% of *de novo* mutations likely occur as PMMs in children. Unexpectedly, we found a significant burden of synonymous PMMs in probands that are predicted to alter splicing. We found no evidence of missense PMM burden in the full cohort. However, we did observe increased signal for missense PMMs in families without germline mutations in probands, which strengthens in genes intolerant to mutations. We also determined that 7-11% of parental mosaics are transmitted to children. Parental mosaic mutations make up 6.8% of all mutations newly germline in children, which has important implications for recurrence risk. PMMs intersect previously implicated high confidence and other ASD candidate risk genes, further suggesting that this class of mutations contribute to ASD risk. We also identified PMMs in novel candidate risk genes involved with chromatin remodeling or neurodevelopment. We estimate that PMMs contribute risk to 4-8% of simplex ASD cases. Overall, these findings argue for future studies of PMMs in ASD and related-disorders.

## Introduction

Autism spectrum disorder (ASD [MIM: 209850]) has a strong genetic component and complex genetic architecture. Over the past decade, technological advances have allowed for the genomewide discovery of rare inherited and *de novo* mutations in ASD cohorts, including: copy number variants (CNVs), structural variants, single nucleotide variants (SNVs), and small insertions and deletions (indels).^1–13^ These studies, especially those focused on simplex cohorts (single affected individual within a family), have revealed a strong burden of *de novo* mutation that implicates hundreds of independent loci in ASD risk as well as, to a lesser yet significant extent, rare inherited mutations. Moreover, while many novel high confidence risk loci and genes have emerged from these studies, the full complement of risk factors and mechanisms have yet to be fully elucidated.

Postzygotic mutations occur after fertilization of the embryo. Depending on their timing and cell lineage, these mutations may be found in the soma, resulting in somatic mosaicism, or the germ cells, resulting in gonadal mosaicism. Mutations occurring during early embryonic development can result in both types of mosaicism.^14^ Gonadal mosaicism is usually only identified when a mutation is transmitted to multiple offspring. If the gonadal mutation cannot be detected in peripheral tissues of the parent, this argues for a germline only origin. For simplicity, we will refer to these mutations generally as postzygotic mosaic mutations (PMMs), as in most cases their contribution to the germline is unknown. PMMs accumulate over an individual’s lifetime and have been shown to have a similar mutation spectrum to germline *de novo* mutations (GDMs).^15^ In addition to the well-known role of somatic mutations in cancer, PMMs have been firmly implicated in several neurodevelopmental/brain disorders including epilepsy, cortical malformations, RASopathies, and overgrowth syndromes.^16–22^ Pathways underlying some of these syndromes, e.g. PI3K/ATK/mTOR and RAS-MAPK, are also implicated in syndromic and nonsyndromic ASD. The mosaic nature of these mutations can make them difficult to identify with current clinical testing, even if the correct gene is known, leading to no diagnosis, misdiagnosis, or misinterpretation of recurrence risk.^17; 23^ Importantly, when and where mutations occur in development can have a dramatic effect on the phenotypic presentation as exemplified by *PIK3CA-related* overgrowth spectrum (PROS).^16; 24^ Moreover, recent data has suggested that even low-level mosaicism (∼1% in affected tissue) can be clinically significant, as shown in the affected skin/brain of Sturge-Weber patients.^25^ Finally, novel genetic etiologies may exist, driven by loci where germline mutations are embryonic lethal.^26^

In previous work focusing on discovering GDMs in simplex ASD families, we were surprised to validate 4.2% of *de novo* mutations as likely mosaic in origin, including nine PMMs and two gonadal mosaic mutations (from a total 260 mutations), suggesting that mosaic mutations might be a common and under-recognized contributor to ASD risk.^2^ A similar observation has been made from *de novo* mutations identified in whole-genome sequencing from simplex intellectual disability (ID) trios.^27^ However, the mutation calling approaches used previously were tuned to detect GDMs. Here, we systematically evaluate the role of PMMs in ASD by leveraging a harmonized dataset^12^ of existing whole-exome sequences (WES) from a well-characterized cohort of ∼2,300 families—the Simons Simplex Collection (SSC), including parents, probands, and unaffected siblings. Our goal was to answer several fundamental questions: 1. What are the rates of PMMs (detectable in whole blood DNA) in parents versus children? 2. How often are mosaic events in the parental generation transmitted to offspring? 3. Do PMMs play a role in ASD risk? 4. Do the targets of GDMs and PMMs overlap?

We first re-evaluated all previously published *de novo* mutations. Using a binomial approach, we found evidence that 11% of SNVs and 26% of indels called with germline methods show allele skewing consistent with mosaicism. We then developed a systematic method for identifying, specifically, SNVs that are likely PMMs from WES (or other next-generation sequencing [NGS] data), which integrates calls from complementary approaches. We benchmarked this approach on a high-coverage WES dataset and then performed extensive validations with molecularly tagged single molecule molecular inversion probes (smMIPs).^28^ Using these validation data, we developed a logistic regression model that is both highly sensitive and specific in distinguishing false positive from true mutations. We also established additional heuristics that when combined with our regression model, generate high confidence mosaic calls. With this more sensitive approach, we recalled genotypes on the SSC cohort and estimate that 22% of *de novo* SNVs are, in fact, PMMs arising in children. Unexpectedly, we found the strongest signal for mutation burden in probands for synonymous PMMs. These synonymous PMMs are enriched for features consistent with potential disruptions of splicing. We did not find strong evidence of missense PMM burden in the full cohort; however, we did observe increasing signal in subsets of the cohort without germline mutations, which is strongest in genes that are intolerant to mutations. We also found strong evidence of transmission of parental mosaic mutations to children. This finding has important potential implications for recurrence risk for families and may explain some instances of parents with subclinical ASD features.^29^ Importantly, we found nonsynonymous (NS) PMMs in high confidence ASD/ID risk genes and other candidate risk genes. We also identify novel candidate risk genes involved with chromatin remodeling or neurodevelopment. Overall, these findings suggest that future studies of PMMs in ASD and related-disorders are warranted. The methods and tools developed here will allow continued discovery of PMMs in future datasets and have potential translational benefits for clinical detection, case management, interventions, and genetic counseling.

## Material and Methods

### Family Selection and Sequence Data

We obtained the initially published^1; 2; 4; 5; 11^ and harmonized reprocessed^12^ WES data from 2,506 families of the Simons Simplex Collection (SSC).^30^ Informed consents were obtained by each SSC recruitment site, in accordance with their local institutional review board (IRB). Oregon Health & Science University IRB approved our study as human subjects exempt as only deidentified data was accessed. Exome libraries were previously generated from whole blood (WB) derived DNA and captured with NimbleGen EZ Exome v2.0 or similar custom reagents (Roche Nimblegen, Inc., Madison, WI) and sequenced using Illumina chemistry (San Diego, CA) at one of three centers: Cold Spring Harbor Laboratory (CSHL), University of Washington (UW), and Yale University School of Medicine. Where individuals had been sequenced by multiple centers, the library with the highest mean coverage was included in the harmonized reprocessed dataset (N. Krumm, personal communication).^12^

For developing our methods, we initially selected 24 family quads (the *pilot 24*) that had WES independently performed by all three centers.^11^ WES data were merged and then reprocessed to match the harmonized dataset.^12^ We then expanded to a cohort of 400 additional independent quad families (the *pilot 400*) with high median WES coverages,^11^ also requiring proportionate distribution across the three centers (Yale: 193, CSHL: 118, UW: 89). Finally, we expanded our analysis to the full SSC harmonized reprocessed dataset.^12^ Families with known identity issues (N. Krumm personal communication) were excluded, yielding 2,366 families, of which 1,781 are quads and 585 are trios (Table S1). One hundred and two families with individuals showing elevated GDM or PMM calls were excluded post variant calling (Supplemental Material and Methods, Figure S1). The cohort used in the downstream analyses included 2,264 families, of which 1,698 are quads and 566 are trios. We removed additional families with low joint coverage depending on the minimum coverage requirement for analyzing variants of different minimum allele fractions (AF) (see Supplemental Material and Methods).

### Evaluating Potential Mosaic Mutations in Previously Published *De Novo* Calls

We combined reported *de novo* mutations for the SSC from previous publications (Table S2).^1; 2; 4; 5; 11; 12^ Allele counts from prior analysis were used where available (N. Krumm, personal communication), and otherwise extracted on a quality-aware basis from mpileups of the corresponding WES using a custom script (*samtools mpileup -B -d 1500 | mPUP -m -q 20 -a count*). Reported mutation calls that had no variant reads from the quality-aware mpileup data were excluded. We focused our analysis on exonic and canonical intronic splice site regions (+/-2 base pairs [bp]). Mutations were considered putative PMMs if significantly skewed from the heterozygosity expectation of 0.5 AF for autosomal and X chromosome sites of females (binomial p <= 0.001). Sex chromosome sites of males were evaluated under a hemizygous expectation. We further analyzed the robustness of the data using additional filters for observed AF (5-35%, 10-35%, 10-25%, or corresponding hemizygous values), or at more strict deviations from the binomial expectation (p <= 0.0001). The observed rates of AF skewed *de novo* mutations were compared with expected null distributions of randomly sampled rare inherited variants by simulation (Supplemental Material and Methods).

### Raw Variant Calling and Annotation

SNVs were recalled on individual samples using VarScan 2.3.2, LoFreq 2.1.1, and our in-house script mPUP (Supplemental Material and Methods). All caller outputs were combined at the individual level and used to generate family-level variant tables. Variants were annotated with ANNOVAR (03/22/15 release [see Web Resources])^31^ against the following databases: RefSeq genes (obtained 2015-12-11), segmental duplications (UCSC track genomicSuperDups, obtained 2015-03-25), repetitive regions (UCSC track simpleRepeat, obtained 2015-03-25), Exome Aggregation Consortium (ExAC) release 0.3 (prepared 2015-11-29), Exome Sequencing Project (ESP) 6500 (prepared 2014-12-22), and 1000 Genomes Phase 3 version 5 (prepared 2014-12-16). Annotation tracks did not include added flanking sequences. Population frequency databases were obtained from the ANNOVAR website. Initially, variants with AFs significantly below 50% (binomial p <= 0.001) were considered putative PMMs. For putative transmitted parental PMMs, which also had skewed AFs in child(ren), we required a significant difference between parent and child AF (Fisher’s exact p <= 0.01), with child AF > parental AF. Only PMM (child or parental) or GDM calls were considered for validation.

### smMIP Design, Capture, and Sequencing

Three to four independent smMIPs were designed against candidate variant sites using the 11-25-14 release of MIPGEN^32^ and a custom in-house selection script (Supplemental Material and Methods). The selected smMIPs were divided into pools with roughly equal numbers (Table S3). Single strand capture probes were prepared similarly to previous approaches with modifications (Supplemental Material and Methods).^32^ DNA samples prepared from WB (entire pilot 24; 78 families pilot 400) and lymphoblastoid cell lines (LCLs) (entire pilot 24) were obtained from the SSC through Rutgers University Cell and DNA Repository (Piscataway, NJ). Probe captures and PCRs to append sequencing adaptors and barcodes were performed as previously described with minor modifications.^33^

Purified capture pools were then combined together for sequencing with NextSeq500 v2 chemistry (Illumina, San Diego, CA). Overlapping reads were merged and aligned using BWA 0.7.1. For each unique smMIP tag, the read with the highest sum of quality scores was selected to serve as the single read for the tag group. Validation outcomes were compared across WB and LCL data (where available) (Table S4).

### Establishing a systematic PMM calling pipeline

We iteratively developed best practices and heuristics through multiple rounds of validation and model development (Supplemental Note: Model Development and Supplemental Material and Methods). We first performed detailed evaluation of the higher depth pilot 24 dataset using smMIPs for validations (Figures S2-S9, Supplemental Note: Model Development and Supplemental Material and Methods). We trained an initial logistic regression model using the pilot 24 initial resolutions, using only calls validated as true PMMs or false positives in the smMIP data. Candidate model predictors were derived from WES data (Supplemental Material and Methods).

We next evaluated pilot 400 quad families (Figures S10-S13). Based on results from the initial validations, for all putative parental transmitted PMMs, we required more significant skew in parental AF (binomial p <= 0.0001), significant difference between parent and child AF (Fisher’s exact p <= 0.01), and child AF > parental AF (Figure S9). All putative PMMs scoring < 0.2 in the initial logistic regression model were excluded. Validations using smMIPs were conducted on calls from 78 of the pilot 400 families. All initial validation positive calls, from both pilot sets, were then subjected to an additional manual review of the WES and smMIP alignments to flag potentially problematic sites prior to modeling.

We trained a refined logistic regression model based on the pilot 400 validation data (Supplemental Material and Methods, Figure S10). We further evaluated this refined model, applying the same filtering parameters as the training set, using the pilot 24 validation calls, which had been selected prior to any modeling or validations.

We evaluated a third set of calls from both pilot sets that had not previously been validated due to data missingness in population frequency datasets (Supplemental Note: Model Development). To better separate germline from mosaic calls based on our empirical validations, we calculated 90% binomial confidence intervals (CI) (Agresti-Coull method) for the variant AFs derived from the WES data using the R *binom* package for all calls. Based upon the distribution of germline resolutions in these data, we re-classified putative PMMs as germline if the upper bound of their observed AF was >= 0.4 (95% CI, one-tailed) (Figure S11). We additionally excluded calls annotated as segmental duplication regions/tandem repeat finder (SD/TRF) sites or mPUP only calls as they had a significantly higher false positive and smMIP probe failure rate (Figure S12). Putative PMMs passing filters from this third set of calls were scored with our refined logistic regression model and excluded from validations if they scored < 0.26. We retroactively applied our refined filtering scheme to all validation calls in order to develop a harmonized set of high confidence resolutions and evaluated sensitivity and PPV of the refined model (Figure S13). Variants with a refined logistic model score >= 0.518 were included for additional analyses.

### Cohort Variant Calling and Burden Analysis

Variants were called from all WES data in the harmonized reprocessed dataset and filtered with our best practice filtering scheme (Supplemental Material and Methods). We additionally required all variants be supported by at least five variant reads and present in no more than two families throughout the cohort to improve PPV for true PMMs (Figure S12). We removed eight remaining variants that had skewed AFs in both the child(ren) and parent. We defined our *high confidence dataset* as those variants with AF >= 5% (based on the AF upper 90% CI) and 45× minimum joint coverage in all family members (Table S5).

For burden analysis, we selected five minimum variant AFs (5%, 7.5%, 10%, 12.5%, 15%) at which to evaluate PMM prevalence across the entire SSC cohort. A variant was included for each subanalysis if its AF upper 90% CI met the minimum AF. For each AF threshold, we determined the minimum total depth (130×, 85×, 65×, 50×, 45×) at which we had approximately 80% binomial probability to observe five or more variant reads (Figure S14). Variants that met minimum coverage requirements in all family members were included in each AF burden analysis and we determined the total number of jointly sequenced bases at or above each depth threshold in each family. Based on these joint coverage values, families in the 5^th^ percentile or lower were excluded; in the 130× analysis the bottom decile was excluded (Figure S15).

To determine mutation burden in the unique autosomal sequence, we first calculated the rate of mutation in each individual by summing all SNVs within a given functional class or gene set, e.g., for missense variants, and dividing by the total number of jointly sequenced bases (diploid, 2n) meeting the minimum coverage thresholds. Rates of mutation were then compared between groups (probands v. siblings, or fathers v. mothers) using, as appropriate, paired or unpaired nonparametric rank tests. To control for multiple comparisons, we used the Benjamini-Yekutieli approach,^34^ which allows for dependent data structures, setting a false discovery rate (FDR) of 0.05. Families of tests were defined based on the dataset and mutation functional class (Supplemental Material and Methods).

We calculated the mean rates for each group by summing all SNVs within a given functional class or gene set and dividing by the total number of jointly sequenced bases (diploid, 2n) for all families meeting the minimum coverage thresholds. Poisson 95% confidence intervals for rates were estimated using the Poisson exact method based on the observed number of SNVs. Means were used for plotting rates and extrapolating variant counts to a full coverage exome.

We performed subcohort burden analyses by separating families on whether or not probands had previously identified GDMs in published call sets. ^1; 2; 4; 5; 11; 12; 35^ Mutations with no read support or flagged as potentially mosaic from our initial analysis of published *de novo* calls were removed (binomial p <= 0.001). We defined two subgroups based on level of disruption The first subgroup included families with probands that have germline *de novo* likely gene disrupting (LGD) SNVs, indels, as well as *de novo* CNVs that affect at least one gene (germline LGD list). The second subgroup included the LGD families and families where probands had any other germline *de novo* NS SNVs or indels (any germline NS list).

To evaluate burden in genes that show evidence of selection against new mutations, we used the recently updated essential gene set^36^, which contains human orthologues of mouse genes associated with lethality in the Mouse Genome Database;^36; 37^ and ExAC intolerant dataset, which denotes the probability of a gene being loss-of-function intolerant.^38^

### Analysis of PMM Properties

The AF distributions between children and parents PMMs were compared by Wilcoxon-rank sum test using the high confidence dataset. To determine the fraction of parental PMMs that may be attributed to lack of grandparental data, we regenerated variant calls from the non-merged reprocessed WES data^12^ for the pilot 24/400 families applying the same refined logistic model and final filters, but ignoring family data. We then fit the observed bimodal AF distributions to normal mixed-models using R package *mixtools*, function *normalmixEM*(), which defined two Gaussian distributions. We separated the calls into two discrete sets. G1 was defined by the mean plus or minus two standard deviations of the leftmost Gaussian model (lower AFs, *μ_1_* = 0.09, *σ_1_* = 0.046). G2 included the remaining higher AF calls. We then determined the fraction of calls remaining in each set after applying transmission filters. We used the fraction of variants retained in the children to estimate the number of variants we expect to remain in the parents if we had sequenced the grandparental generation.

To determine the mutational spectra, we used R package *MutationalPatterns* to extract and plot mutational contexts, calculate their relative frequency within our high confidence GDMs, child PMMs, and parental nontransmitted PMMs.^39^ We downloaded the frequencies of mutation within the 96 possible trinucleotides for each cancer signature (see Web Resources) and used the Pearson method to correlate each signatures mutational spectrum to our own.^40^

Splice site distances for variants were annotated using Variant Effect Predictor (see Web Resources). The absolute value of the shorter of the two distances between donor or acceptor site was chosen as the distance to nearest splice site. To predict the impact on splicing by synonymous variants, we used Human Splice Finder (HSF) version 3.0 and SPANR alpha version (see Web Resources).^41; 42^ For HSF, we used multiple transcript analysis with default settings and results were extracted from HTML format outputs with an in-house script (Table S6). Variants contained within multiple overlapping transcripts with disparate calls were manually filtered based on whether transcripts were coding or had complete stop/start information in the UCSC genome browser (Feb. 2009; GRCh37/hg19). SPANR analysis was performed with default settings and splice altering variants defined as described previously (5% > dPSI percentile or dPSI percentile > 95%).

### Gene Set Enrichment

We examined potential enrichments of missense variants in five different gene sets that have been previously been evaluated using *de novo* mutations,^11^ including an updated version of the essential gene list.^37^ We downloaded genesets from GenPhenF (See Web Resources) and then mapped gene symbols to our RefSeq ANNOVAR annotations. To determine enrichment, we took a similar approach as previously described, using the null length model.^11^ However, we calculated joint coverage for all genes within a set as well as all the genes outside of that set (across the cohort) and used this value to estimate the expected proportion of mutations (*p*). Since more than one gene can overlap any genomic position, for this analysis we counted based on all genes impacted. Thus, if a mutation or genomic position overlapped a gene within the set and outside of the set, it was counted twice. We tested for gene set enrichment using a binomial test in R *binom.test*(*x, n*, *p*), where x = number of genes impact within set, n = total number of genes impacted, *p* = expected mean based on joint coverage.

To determine if genes targeted by missense or synonymous mutations in probands showed enrichment for ASD candidate gene rankings, we used genomewide gene rankings generated from two previous studies.^36; 43^ The *LGD* intolerance ranking is based on the load of LGD mutations observed per gene.^36^ The *LGD-RVIS* is the average rank between LGD and RVIS (another measure of constraint) scores.^36; 44^ *ASD association* rankings are the results of a machine learning approach that uses the connections of ASD candidate genes within a brain-specific interaction network to predict the degree of ASD association for every gene.^43^

### Intersection of PMMs with Previously Published GDMs

Degree of overlap of GDMs and PMMs for different functional classes between probands and siblings was determined using Fisher’s exact test. Both the high confidence and burden (15%-45x) datasets were evaluated. Our high confidence (HC) risk gene set was curated using the 27 ASD genes reported by Iossifov et al. and 65 ASD genes reported Sanders et al. (FDR <= 0.1).^11; 35^ We also included 94 genes enriched for GDMs in developmental disorders from the Deciphering Developmental Disorders study and 91 genes from the Autism Spectrum/Intellectual Disability (ASID) network study.^45; 46^ Combined the HC risk gene sets includes 171 unique genes.

## Results

### Reanalysis of Previously Reported *De Novo* Mutations

We began by analyzing the existing set of previously reported exonic or canonical intronic splice site *de novo* mutations in the SSC (Table S2).^1; 2; 4; 5; 11; 12^ We evaluated 5,076 SNVs (probands: 2,996; siblings: 2,080) and 416 small indels (probands: 273; siblings: 143) (Figures 1A-D and S16-19 and Tables S7-S12). Variants had a mean depth of 77.5×. We found an excess of mutations with observed AFs lower than expected for germline events using a binomial threshold of 0.001 (Figures 1A, C and Table S7). We evaluated the likelihood of this excess within the autosomes sequence by simulating a null distribution from rare inherited SNVs (Supplemental Materials and Methods, Figures 1B, S17 and Table S8). For autosomal *de novo* SNVs, we observed that 305/2,893 (11%) of affected proband calls and 191/1,993 (10%) of unaffected sibling calls show evidence of being PMMs. For rare inherited SNVs, we never observed the same degree of skewing of calls with lower AFs (simulation means: probands-80/2,893 [2.8%]; siblings-56/1,993 [2.9%]; p < 0.0001, by simulation). A higher potential PMM rate is observed in sites that annotated as SD/TRF loci, 55/231 (24%) in probands and 28/144 (20%) in siblings (p = 0.0166 and 0.41, respectively, by simulation). These SD/TRF sites are known to be more prone to false PMM calls due to uncertain mapping of WES reads. However, these SD/TRF loci only represent 9% of the called mutations and thus have a modest effect on the overall rate. We observed a similar rate of potential SNV PMMs (8-9%) when applying a range of additional AF cutoffs (5-35%, 10-35%, 10-25%), more strict binomial deviations (p <= 0.0001), or both (Table S9 and Table 10), suggesting these are robust estimates. In sharp contrast, we did not observe an excess of calls with higher than expected AFs (Table S8). In probands, the relative proportion of mutation types, i.e. fraction of synonymous or missense, were similar between calls classified as GDMs versus PMMs. Interestingly, in siblings, the fraction of synonymous calls appears reduced in PMMs compared with GDMs (48/217 [22%] v. 531/1,863 [29%], p = 0.054, two-sided Fisher’s exact) (Table S7).

For indels, we also observed a large number of potential PMMs exceeding the binomial expectation (Figures 1B,D and Table S11), with more variability overall between probands and siblings (57/268 [22%] v. 48/140 [35%], respectively, p = 0.005, two-sided Fisher’s exact). For rare inherited indels, we never observed the same degree of skewing of calls with lower AFs (simulation means: probands-16/268 [6%]; siblings-10/140 [7%]; p <0.0001, by simulation) (Figures 1D, S18-S19 and Table S8). Similar to SNVs, we found an elevation in the rate for SD/TRF loci (probands-7/18 [39%]; siblings-9/16 [56%]; p = 0.0003 and <0.0001, respectively, by simulation). However, the percent PMM estimates were less robust, compared with SNVs, when applying additional AF cutoffs, more strict binomial deviations, or both (Tables S11 and S12). For example, the overall PMM rates using the stricter binomial threshold reduced to 40/268 (15%) for probands and 33/140 (24%) for siblings (p = 0.045, two-sided Fisher’s exact), which nevertheless still exceeded the null expectation (p < 0.0001, by simulation) (Table S8). We observed no *de novo* indels with significantly deviated higher AFs.

We next examined validation data previously reported or available for a subset (63/545) of the predicted mosaic calls, which included Sanger and NGS data. We found 39/63 (62%) calls showed strong evidence of allele skewing in the validation data (Table S2). These data argue the majority of these calls are *bona fide* PMMs but that systematic approaches tuned to detecting PMMs are still needed.

### Developing a Systematic Mutation Calling Framework

We sought to perform a systematic analysis of PMMs with methods specifically geared toward SNV mosaic mutations, which do not require a matched ‘normal’ tissue data comparison (Figure 1E). Moreover, we expected a large number of suspected PMM calls to be false because of random sampling biases, mapping artifacts, or systematic sequencing errors. Therefore, we worked to build a robust calling framework that would integrate different approaches and could be empirically tuned based on validation data.

**Figure 1.**
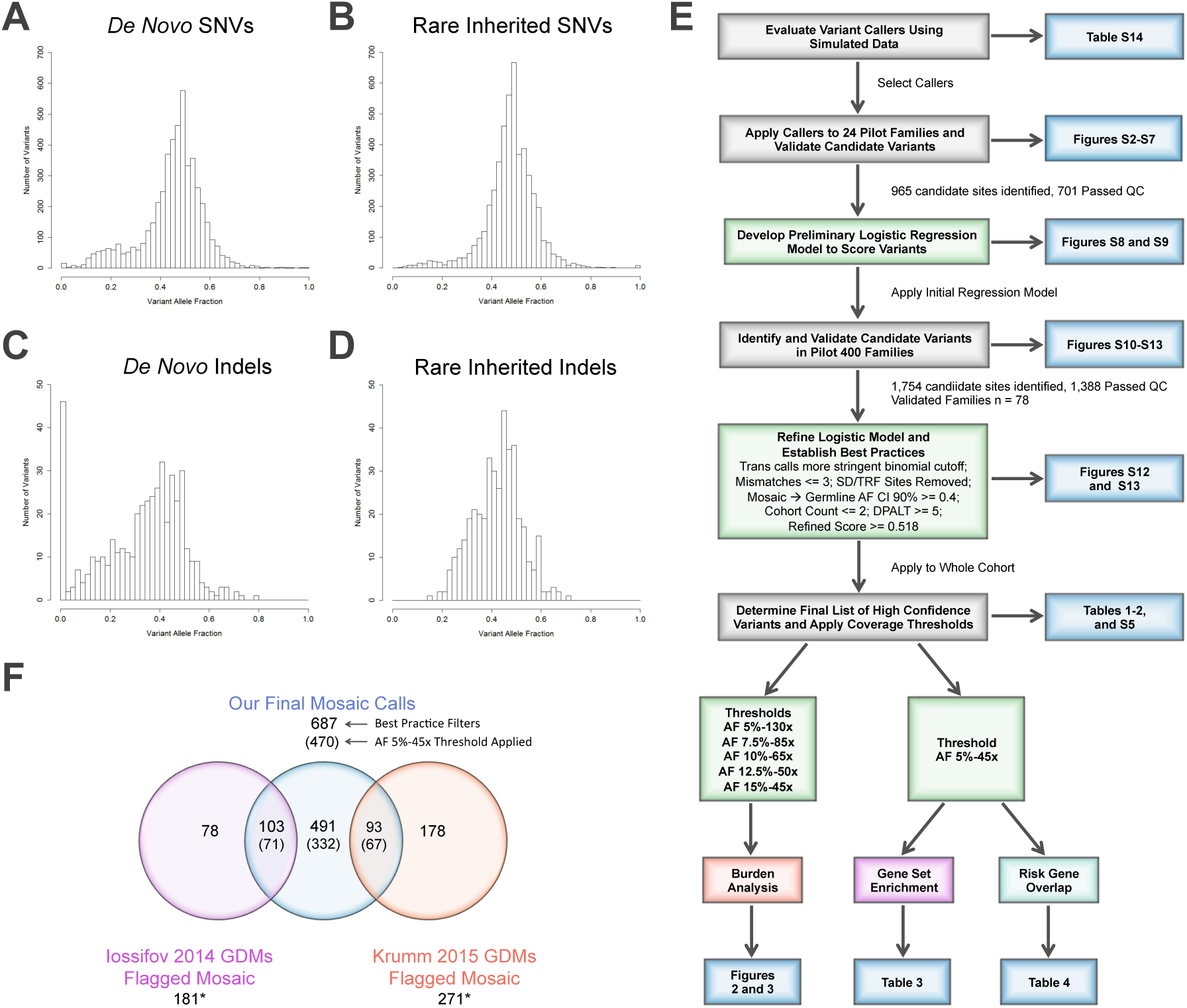
Re-Evaluation *De Novo* Mutations in the Simons Simplex Collection (SSC) (A-D) Histograms showing the allele fraction distributions of previously published *de novo* or rare inherited mutations in the SSC.
(A) Published *de novo* SNVs show an elevated number of low allele fraction calls that are potentially PMMs (left tail).
(B) Representative histogram from a random sampling of published rare inherited SNVs. The number of low allele fraction calls is substantially fewer compared to *de novo* SNVs (left tail).
(C) Published *de novo* indels show an elevated number of low allele fraction calls (left tail) that are potentially PMMs as well as an overall shifted distribution.
(D) Representative histogram from a random sampling of published rare inherited indels. Similar to SNVs, the number of low allele fraction calls is substantially fewer compared to *de novo* indels (left tail).
(E) Schematic showing an overview of our systematic approach to developing a robust PMM calling pipeline and applying it to the SSC. Key analyses and display items are indicated. Abbreviations: Trans calls-calls showing evidence of transmission from parent to child, SD/TRF-segmental duplications/tandem repeats, AF-allele fraction, CI-confidence interval, and DPALT-Q20 alternative allele depth.
(F)Venn diagram showing the intersection of previously published *de novo* mutations initially flagged as potentially PMMs (binomial p <= 0.001) and our PMM calls after applying final filters. Numbers in parentheses are calls remaining after applying an AF 5%-45x joint coverage threshold. *Our pipeline identified an additional 37 calls (29 from Iossifov et al.^11^ and 8 from Krumm et al.^12^), which overlapped the published calls flagged as potentially mosaic, but were re-classified as likely germline based on their AF CIs. Note: Krumm 2015 dataset only reported newly identified calls and therefore does not intersect the Iossifov 2014 dataset.

We evaluated several standalone (single sample) SNV mosaic mutation callers, including Altas2^47^, LoFreq^48^, Varscan2^49^, and a custom read parser (mPUP) using simulated data containing artificial variants at 202 loci. We found that within the simulated data, caller sensitivity greatly varied at different depths and AFs, but many had high PPV (Table S13 and Table S14). Based on their complementary performances at different depths and AFs, we selected Varscan2, LoFreq, and mPUP for further evaluation.

Next, we took advantage of the fact that 24 quad families (96 individuals) had WES independently generated by three centers, providing an opportunity to empirically evaluate these methods on a combined remapped and merged high-depth WES dataset (merged pilot 24: average mean coverage 208x) (Figures S2B and S15A). We obtained high confidence validation data from at least one DNA source using smMIPs and Illumina sequencing for 645/902 (72%) of the predicted PMM and 56/63 (84%) of the GDM sites (Figure S3 and Table S4). Not surprisingly, we found the majority of the PMMs predicted by a single variant caller were false positives (345/347, 99%), whereas those called by at least two other approaches had a better PPV (162/298, 54%) (Figure S7). In addition, a number of PMMs were in *cis* with existing heterozygous polymorphisms. PMM alleles tracked with specific haplotypes, but were absent from a number of overlapping reads, strongly suggesting that these are *bona fide* postzygotic events (Figure S4). We further found that for transmitted variants, we could eliminate most of the mischaracterized calls that validated as parental germline by requiring a more significant binomial deviation and performing a Fisher’s exact test of the read counts from the parent-child pair (Figure S9). Some of these transmitted variants showed consistently skewed AFs that transmitted in a Mendelian fashion, suggesting they are either systematically biased or multicopy sites that we co-sampled (Figure S5).

Using these pilot 24 validation data, we constructed an initial logistic regression model (Supplemental Material and Methods). We then applied this initial logistic regression model and additional filters for ambiguous transmitted sites to an independent set of 400 quad families (Material and Methods, Figure S10). We performed smMIPs validation on WB DNA samples from 78 of these quads and obtained high confidence validation data on 1,388/1,754 sites.

Based on manual inspection of the WES and smMIP alignment data, we identified additional features associated with poor prediction outcomes or problematic genomic regions, including multiple mismatches within the variant reads and presence in multiple families (Figures S6 and S12A-B). We added filters based on these features to the pilot 400 validation set and built a refined logistic regression model (Figure S10). The model performed well in 3-way cross validations with sensitivity estimated at 92% and PPV at 80% (threshold 0.26) (Figure S13A). To further evaluate this model, we rescored the pilot 24 validation sites with and without additional filters (Material and Methods). Importantly, these calls were selected and validated prior to model development, giving an independent set of data to evaluate performance. These data performed better than the training data (after removing mPUP only calls), likely due to the increased WES coverage of the pilot 24 samples with sensitivity of 94% and PPV of 83% (threshold 0.26) (Figures S13C-D).

As we specifically developed our model to separate predicted PMM calls that validate as false versus true variants, regardless of whether they were mosaic or germline, we subsequently examined the validation data to determine if an additional heuristic could further distinguish true mosaic calls from calls that validated as germline. We observed that calls validating germline tended to have higher observed WES AFs. We calculated the 90% binomial CI (95% one-sided) for the observed AF as a potential complement to the observed significant binomial deviations. We found that the vast majority 112/113 (99%) of validated PMM calls had upper CI bounds that remained below 0.4, while bounds for the majority of true germline calls 25/33 (76%) fell above this threshold (Figure S11). In addition, we observed that a significant fraction of the false positive calls exceeding our logistic score threshold (5/26, [19%]) were annotated as SD or TRF sites (Figures S12C-D). Moving forward we chose to remove these SD/TRF sites and re-classify mosaic versus germline status based on the AF binomial CI.

Finally, we scored and filtered the pilot cohorts using these parameters and conducted a third set of validations on PMM and GDM calls not previously evaluated (Supplemental Note: Model Development). We evaluated these new data and our previous validation calls under these harmonized filters (Figures S13E-F). We observed that across all test sets (excluding training data), both sensitivity and PPV converged at a logistic score of 0.518 (sensitivity 0.83, PPV 0.85). At this score threshold, 21/22 (95%) of mosaic predictions that validated as true variants were confirmed as mosaic in children (all test sets). We chose to use this more stringent score threshold for our subsequent burden analysis. In addition, we removed calls with less than five variant allele reads as these disproportionately contributed to false calls (Figure S12E).

### Evaluation of Mutation Rates and Burden in Children with ASD

Using this approach we recalled SNVs in the SSC, in both children and parents, from the existing harmonized reprocessed WES data (average mean coverage 89x).^12^ We identified 687 total PMMs originating in the children from 1,699 quads and 567 trios passing SNV QC metrics (Tables 1 and S5). We re-identified 3,445/4,198 previously published SNVs GDMs, which were not flagged as potentially mosaic, and 1,064 novel calls. Applying our high confidence call set criteria (5% minimum AF and 45x joint coverage) resulted in 470 PMMs, of which 332 were not part of the published GDM calls (Figure 1F and Table 1). Of the 452 previously published SNV GDMs that we initially flagged as potentially mosaic, 233 were called by our approach (196 as mosaic), of which 157 remained in our high confidence call set (138 as mosaic, 19 as reclassified germline) (Figure 1F). Likewise, applying the high confidence call set criteria reduced the GDM count to 1,677, of which only 10 were novel. Compared to our analysis of previously published *de novo* SNVs, we observed a higher fraction of mosaic mutations amongst the *de novo* calls in children, 470/1,677 (22%), consistent with increased sensitivity of our mosaic targeted approach (Table S15).

**Table 1.** PMM Counts in Children Across Different Allele Fraction and Coverage Thresholds

|  |  | syn | mis | non+splice | Total |
| --- | --- | --- | --- | --- | --- |
| Best Practice Filters |  |  |  |  |  |
| Quads | Pro | 94 | 194 | 20 | 308 |
|  | Sib | 62 | 203 | 15 | 280 |
| Trios | Pro | 26 | 63 | 6 | 95 |
|  | Total Pro | 120 | 257 | 26 | 403 |
| AF 5%-45x High Confidence |  |  |  |  |  |
| Quads | Pro | 58 | 131 | 12 | 201 |
|  | Sib | 42 | 133 | 10 | 185 |
| Trios | Pro | 22 | 53 | 6 | 81 |
|  | Total Pro | 80 | 184 | 18 | 282 |
| AF 15%-45x Burden* |  |  |  |  |  |
| Quads | Pro | 24 | 65 | 5 | 94 |
|  | Sib | 20 | 66 | 5 | 91 |
| Jointly Covered Bases: 24.5 |  |  |  |  |  |
| Trios | Pro | 8 | 30 | 0 | 38 |
|  | Total Pro | 32 | 95 | 5 | 132 |
| Jointly Covered Bases: 9.7 |  |  |  |  |  |
| AF 12.5%-50x Burden* |  |  |  |  |  |
| Quads | Pro | 32 | 67 | 5 | 104 |
|  | Sib | 16 | 80 | 6 | 102 |
| Jointly Covered Bases: 22.3 |  |  |  |  |  |
| Trios | Pro | 12 | 31 | 2 | 45 |
|  | Total Pro | 44 | 98 | 7 | 149 |
| Jointly Covered Bases: 8.9 |  |  |  |  |  |
| AF 10%-65x Burden* |  |  |  |  |  |
| Quads | Pro | 38 | 63 | 6 | 107 |
|  | Sib | 20 | 76 | 4 | 100 |
| Jointly Covered Bases: 16.7 |  |  |  |  |  |
| Trios | Pro | 12 | 31 | 1 | 44 |
|  | Total Pro | 50 | 94 | 7 | 151 |
| Jointly Covered Bases: 6.8 |  |  |  |  |  |
| AF 7.5%-85x Burden* |  |  |  |  |  |
| Quads | Pro | 31 | 56 | 6 | 93 |
|  | Sib | 18 | 66 | 5 | 89 |
| Jointly Covered Bases: 11.4 |  |  |  |  |  |
| Trios | Pro | 11 | 28 | 4 | 43 |
|  | Total Pro | 42 | 84 | 10 | 136 |
| Jointly Covered Bases: 4.7 |  |  |  |  |  |
| AF 5%-130x Burden* |  |  |  |  |  |
| Quads | Pro | 20 | 35 | 4 | 59 |
|  | Sib | 12 | 35 | 2 | 49 |
| Jointly Covered Bases: 5.1 |  |  |  |  |  |
| Trios | Pro | 10 | 18 | 5 | 33 |
|  | Total Pro | 30 | 53 | 9 | 92 |
| Jointly Covered Bases: 2.0 |  |  |  |  |  |

Abbreviations: AF-allele fraction, Pro-proband, Sib-sibling, syn-synonymous, mis-missense, non + splice-nonsense and canonical splicing. Bases in billions. \*PMMs in sex chromosomes were excluded in this set.

In the SSC, *de novo* large gene disrupting CNVs, likely gene disrupting (LGD) GDMs, and missense GDMs have been shown to have a greater mutation burden in individuals affected with ASD versus their unaffected siblings.^4; 11; 50^ We reasoned that the burden of PMMs might differ based on embryonic timing given that an early embryonic mutation would contribute more substantially to postembryonic tissues. Therefore, we evaluated burden across the entire SSC cohort at several defined minimum AFs, as a surrogate for embryonic time, and corresponding joint family coverage thresholds (AF-COV): 5%-130×, 7.5%-85×, 10%-65×, 12.5%-50×, and 15%-45× (Figure S14 and Table 1).

We first examined the mutation burden of the unique autosomal coding regions in quad families exclusively as they provided a matched set of child samples (Material and Methods). Within our 15%-45× GDM calls, we recapitulated the previously observed mutation burdens for missense (p = 0.003, one-sided Wilcoxon signed-rank test (WSRT)) and nonsense/splice mutations (p = 0.00025, one-sided WSRT) and lack of burden for synonymous mutations, demonstrating these findings are robust to removing potential PMM calls. Mean rates were similar across minimum AF-COV thresholds with a slight trend toward lower rates at high minimum depths. In contrast, for PMMs we see an increase in mutation rate at higher depths and corresponding lower AFs. This is in line with expectations as newer mutations (lower AFs) would accumulate during development. Given the low number of nonsense/splice mutations (Figure 2A), we restricted our burden analyses to synonymous and missense PMMs. We observed no bulk mutation burden signal for missense PMMs (Figure 2B). Unexpectedly, we observed an increased burden of synonymous PMMs in probands (Figure 2C). The signal was strongest in the 12.5%-50× subanalysis with probands having twice as many mutations (32 in probands or 7.2×10^−10^/base pair v. 16 in siblings or 3.6×10^−10^/base pair, p = 0.0024, two-sided WSRT, FDR < 0.05). This trend continued for the three lower AF windows, but these did not exceed an FDR of 0.05. We extrapolated the observed mean per base rates to the full unique autosomal RefSeq exome (31,854,496 bases/haplotype, including canonical splice sites) in order to calculate the average differential between probands and siblings, similar to the analysis performed previously for GDMs.^11^ Based on the 12.5%-50× data, we found probands had a rate of 0.046 synonymous PMMs per exome and siblings 0.023, suggesting 50% of proband synonymous PMMs contribute to ASD risk. The differential between probands and siblings was 0.023, which translates to 2.3% of simplex cases in the overall cohort harboring a synonymous PMM related to ASD risk.

**Figure 2.**
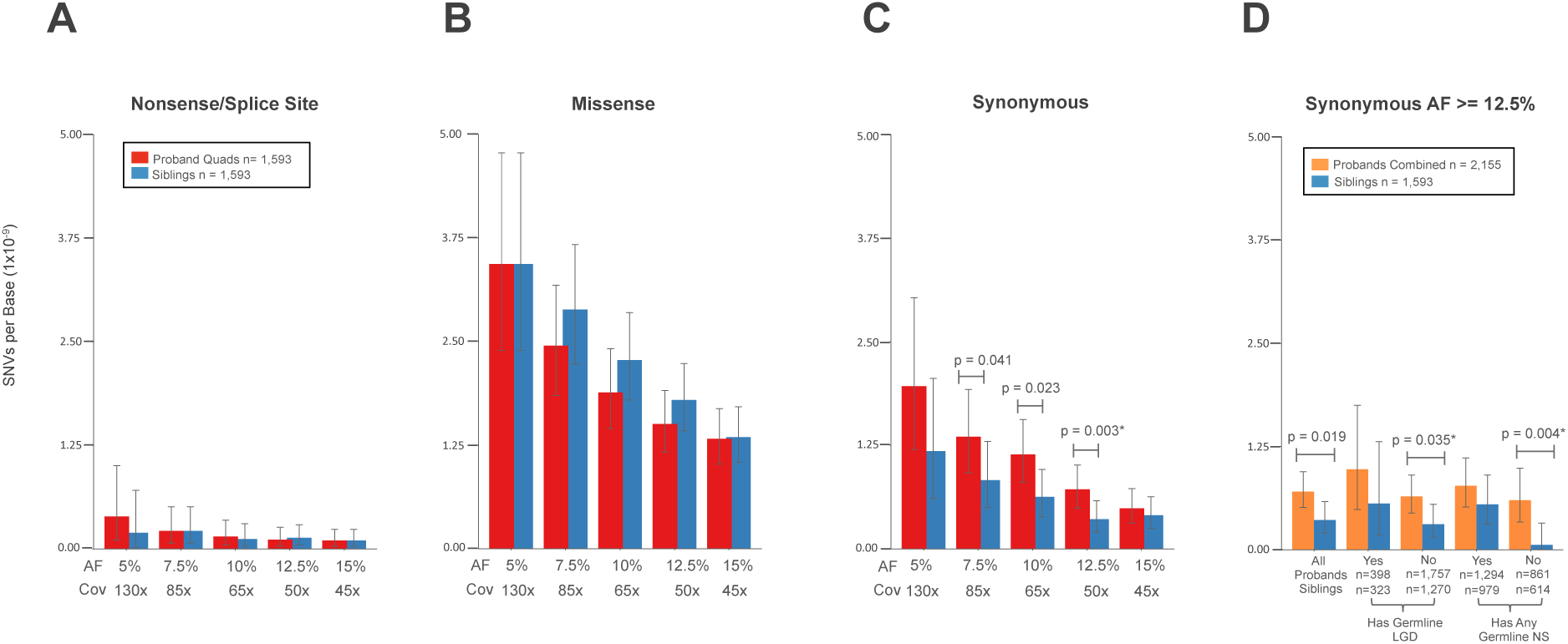
Rates and Burden of SNV PMMs in the Simons Simplex Collection (SSC) (A-C) Rates and burden analyses of PMMs in quad families of the SSC. Mean rates with 95% Poisson CIs (exact method) are shown.
(A) Nonsense/splice PMM rates are similar and not evaluated further given their low frequency.
(B) Missense PMMs show no evidence of burden in probands from quad families.
(C) Synonymous PMMs show an unexpected burden in probands from quad families. Significance determined using a two-sided Wilcoxon signed-rank test. *FDR < 0.05 using the Benjamini-Yekutieli approach.
(D) Analysis of synonymous PMMs at AF 12.5%-50x in the full SSC and subcohorts. Mean rates with 95% Poisson CIs (exact method) are shown for combined probands (quad + trio families) and unaffected siblings. SSC subcohorts: All-all families within the cohort passing quality criteria; Has Germline LGD-denotes whether or not proband in family has a LGD GDM or gene disrupting *de novo* CNV; Has Any Germline NS-denotes whether or not proband in family has any NS GDM (includes the LGD set). Significance determined using a two-sided Wilcoxon rank sum test. *FDR < 0.05 using the Benjamini-Yekutieli approach.

We next combined the data from quad and trio-only (father, mother, proband) families to increase the number of mutations and conducted an exploratory analysis of mutation rates in subsets of the full cohort. Since a large fraction of the SSC has GDM events that are likely contributory, we reasoned that grouping families by presence or absence of proband GDMs of different severity (LGD/disruptive CNV v. any NS) might improve our ability to detect any PMM signal that might be present. Based on the 12.5%-50× data in families without a germline LGD, we observed burden signal similar to the full cohort (p = 0.004, two-sided Wilcoxon rank-sum test [WRST], FDR <0.05). However, the full cohort data did not meet the FDR threshold using the less powerful unpaired test data. In contrast, for the families without any reported NS GDMs, we observed a dramatic depletion of synonymous PMM events in the unaffected siblings, with a proband to sibling rate ratio of 10 (Figure 2D). In this group without NS GDMs, this equates to 0.038 synonymous PMM events per proband exome and 0.004 per sibling exome (differential of 0.034), suggesting 89% of this mutation class contributes to ASD risk.

Next, we examined missense PMMs using the two cohort subgroupings at the 15%-45× threshold. We observed a non-significant trend toward burden of missense PMMs in probands for families either without any LGD GDMs (rate ratio 1.28) or without any NS GDMs (rate ratio 1.49) (p = 0.085 and p = 0.076, respectively, one-sided WRST) (Figure 3A). It has now been well documented using several approaches that LGD GDMs in probands show enrichments for genes that are highly conserved/intolerant to LGD mutations.^11; 50; 51^ We reasoned that missense PMMs relating to ASD risk could also show similar enrichments. We selected two intolerant gene sets, an updated set of essential genes (n = 2,455)^37^ and the recently published ExAC intolerant set (n = 3,232).^38^ These subanalyses showed increased effect sizes, but none of these results exceeded a FDR of 0.05. For both essential and ExAC intolerant sets, we observed similar trends for enrichments of missense PMMs in probands (rate ratios 1.4, p = 0.093 and p = 0.13, respectively, one-sided WRST).

**Figure 3.**
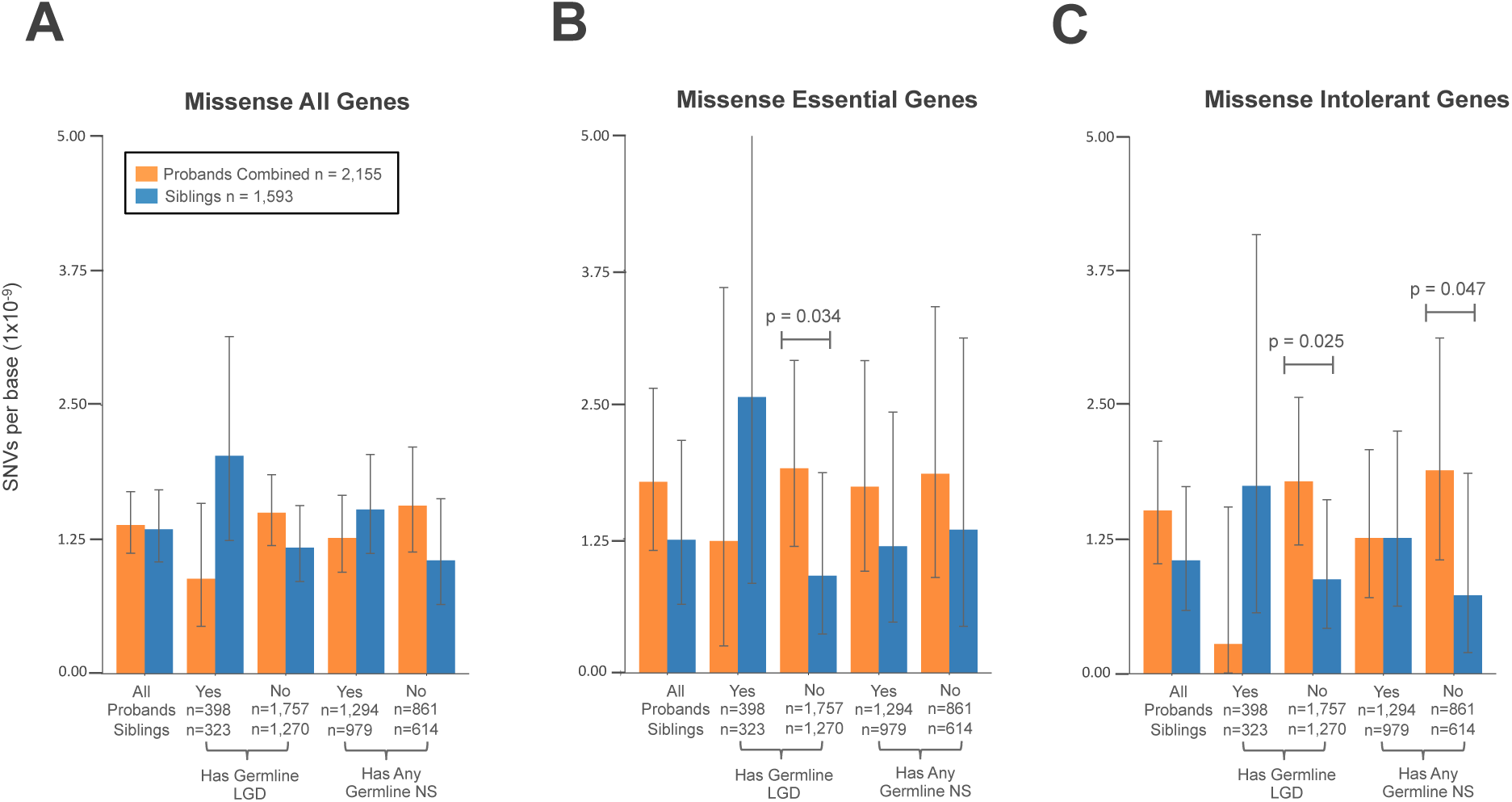
Rates and Burden of Missense PMMs in Subcohorts and Gene Sets. For all plots, the 15%-45× burden call set was used Mean rates with 95% Poisson CIs (exact method) are shown. SSC subcohorts: All-all families within the cohort passing quality criteria; Has Germline LGD-denotes whether or not proband in family has a LGD GDM or gene disrupting *de novo* CNV; Has Any Germline NS-denotes whether or not proband in family has any NS GDM (includes the LGD set). Significance determined using a one-sided Wilcoxon rank sum test. No comparisons met a FDR < 0.05 using the Benjamini-Yekutieli approach.

A. Splitting by subcohort shows trends for increased missense PMM burden in families where probands do not have reported germline mutations.
B. Evaluating mutations specific for the essential gene set shows stronger proband burden in the without any germline LGD subcohort.
C. Similarly, evaluating mutations specific for the intolerant gene set shows stronger proband burden in the without any germline LGD or without any germline NS subcohorts.

When combining these two approaches, which subdivide the cohort and gene targets, we saw the strongest effects. In the subset of families without LGD GDMs, we saw a stronger effect for both essential and ExAC intolerant genes (rate ratios 1.4 and 2, p = 0.034 and p = 0.025, respectively, one-sided WRST). We observed similar results when restricting to quad only families. Missense PMMs in essential/intolerant genes occur at a rate of 0.12 events per exome in probands who do not have a LGD GDM and at a rate of 0.11 for intolerant genes (0.06 and 0.05 for siblings, respectively, differentials 0.06). Interestingly, the families without any NS GDMs showed the largest effect in the ExAC intolerant set (ratio 2.6, p = 0.047, one-sided WRST), but similar rates to the full cohort in the essential gene set (ExAC: 0.12 events per proband, 0.05 per sibling, 0.07 differential). Based on these differentials, we estimate that 6-7% of probands without LGD or NS GDMs have a missense PMM in an essential/intolerant gene contributing to risk. Adjusting to the full cohort, gives a range of 2.1-5.6% of probands harboring a missense PMM in an essential/intolerant gene related to ASD risk.

### Parental PMM Rates and Transmission

We divided PMMs originating in the SSC parents into transmitted and nontransmitted mutations (Tables 2 and S15 and Figure S4). We identified 1,293 nontransmitted (654 in fathers and 639 in mothers) and 92 transmitted (52 in fathers and 40 in mothers) total PMMs in our high confidence call set. For transmitted mutations, which by definition require the postzygotic mutation contribute to both soma and germline, we required a stricter deviation from the binomial expectation based on empirical validation data (p <= 0.0001). The overall PMM rates were similar between fathers and mothers for different mutation functional classes and across AF-COV windows (Figure S20). We did see a noticeable, though non-significant, trend toward higher transmitted synonymous rates in fathers. As with the children’s PMMs, we observed slight increases in mutation rate at higher depths and corresponding lower AF thresholds. In the high confidence call set, we found the PMM rate to be 2.6-fold greater in the SSC parents relative to their children. However, we expect that some of this bias may be due to differences in filtering out transmitted sites that show false mosaic signal, as we do not have the previous generation, i.e. grandparents, to compare to as we do for the children. Therefore, we looked at variants in a subset of the cohort and determined the fraction of variants remaining in children before and after applying transmission filters. Using this rate, we estimated the number of PMMs expected to be filtered from the parental calls based on transmission. We estimate that 40% of our parental PMM calls are in excess of what is expected and likely attributed to incomplete filtering (Figure S21). Applying this correction reduces the parental excess PMM rate to 1.6-fold greater. Based on the children, two-thirds of filtered calls appear to be systematically biased as they are skewed in both generations. One-third of these calls are only skewed in a single generation with AFs > 20%, suggesting they are likely stochastic events.

The increased rate of PMMs in parents compared to children is in line with previous observations that PMMs accumulate with age.^15; 52^ We also observed an overall trend toward an increase in the rate of PMMs with parental age for both sexes (Figure S22A). The rate of PMMs markedly increases after age 45 and there is a significant difference in rate between parents younger than 45 as compared to those 45 and older (mothers-rate ratio 1.2, p = 0.04; fathers-rate ratio 1.3, p = 0.01, one-sided WRST) (Figure S22B). We also saw that the number of individuals with multiple PMMs (adjusted for coverage differences) within a given age range increased as well (Figure S22C). Recent studies have also demonstrated a rise in PMMs in particular genes that result in aberrant clonal expansions (ACEs) that are specific to blood cells.^52–55^ We did not find strong evidence for enrichment of PMMs in 42 genes with recurrent ACE associated mutations from three studies of hematopoietic clonal expansion (parents-obs: 9, exp: 6.6, p = 0.17; children-obs: 5, exp: 2.3, p = 0.07; two-sided binomial).^53–55^ However, among the parents we did find recurrent PMMs in two of the most frequently mutated ACE-related genes, *DMNT3A* (four nonsense and one missense) and *TET2* (two missense). These PMMs did occur in relatively older individuals for our cohort, 45-50 years old. Two missense PMMs in *TET2* were also observed in the children.

Within the 45× joint coverage data, we found that 7-10% of parental PMMs were transmitted to one or more children depending on the minimum AF threshold (5% v. 15%), which requires an early embryonic origin (Table 2). Moreover, in our high-depth validation data with final filters applied, we found 1/164 GDM predictions showed evidence of low AF in parental DNA, which was not detected by WES (Table S4). Taken together, these data argue that at a minimum 7-11% of new mutations are likely occurring prior to germline specification. In addition to these mutations that showed evidence of mosaicism in the parental WES/smMIP data, we also identified six obligate mosaics given their *de novo* presence in two offspring, i.e. gonadal mosaic mutations (Table S5). These gonadal mutations make up 0.66% of the germline mutations in the children. Finally, within the quad families of our high confidence call set we did observe skewing of transmission to siblings (18 to both, 39 siblings, 22 probands).

**Table 2.** PMM Counts in Parents Across Different Allele Fraction Coverage Thresholds

|  |  | syn | mis | non+splice | Total |
| --- | --- | --- | --- | --- | --- |
| **Best Practice Filters** |  |  |  |  |  |
| Nontrans | Fa | 259 | 543 | 54 | 857 |
|  | Mo | 266 | 570 | 41 | 878 |
| Trans | Fa | 21 | 41 | 1 | 63 |
|  | Mo | 12 | 37 | 0 | 49 |
| **AF 5%-45x High Confidence** |  |  |  |  |  |
| Nontrans | Fa | 196 | 418 | 40 | 654 |
|  | Mo | 199 | 405 | 35 | 639 |
| Trans | Fa | 19 | 32 | 1 | 52 |
|  | Mo | 7 | 33 | 0 | 40 |
| **AF 15%-45x Burden\*** |  |  |  |  |  |
| Nontrans | Fa | 114 | 261 | 19 | 394 |
|  | Mo | 130 | 267 | 15 | 412 |
| Trans | Fa | 19 | 32 | 1 | 52 |
|  | Mo | 6 | 31 | 0 | 37 |
| Jointly Covered Bases: 34.2 |  |  |  |  |  |
| **AF 12.5%-50x Burden\*** |  |  |  |  |  |
| Nontrans | Fa | 126 | 276 | 22 | 424 |
|  | Mo | 130 | 281 | 18 | 429 |
| Trans | Fa | 16 | 30 | 1 | 47 |
|  | Mo | 6 | 30 | 0 | 36 |
| Jointly Covered Bases: 31.2 |  |  |  |  |  |
| **AF 10%-65x Burden\*** |  |  |  |  |  |
| Nontrans | Fa | 121 | 229 | 18 | 368 |
|  | Mo | 110 | 229 | 19 | 358 |
| Trans | Fa | 11 | 23 | 1 | 35 |
|  | Mo | 4 | 20 | 0 | 24 |
| Jointly Covered Bases: 16.7 |  |  |  |  |  |
| **AF 7.5%-85x Burden\*** |  |  |  |  |  |
| Nontrans | Fa | 90 | 177 | 19 | 286 |
|  | Mo | 92 | 180 | 19 | 291 |
| Trans | Fa | 5 | 15 | 1 | 21 |
|  | Mo | 2 | 13 | 0 | 15 |
| Jointly Covered Bases: 16.1 |  |  |  |  |  |
| **AF 5%-130x Burden\*** |  |  |  |  |  |
| Nontrans | Fa | 53 | 110 | 15 | 178 |
|  | Mo | 49 | 101 | 9 | 159 |
| Trans | Fa | 3 | 4 | 0 | 7 |
|  | Mo | 1 | 5 | 0 | 6 |

Abbreviations: AF-allele fraction, FA-father, Mo-mother, syn-synonymous, mis-missense, non + splice-nonsense and canonical splicing. Bases in billions. \*PMMs in sex chromosomes were excluded in this set.

### Properties of PMMs

We next examined additional properties of PMMs from the high confidence call set (Table S5). We examined the AF distributions of PMMs and found that between parents (fathers and mothers), and likewise between children (probands and siblings), AF distributions were similar (Figure 4). Therefore, we combined parental calls and child calls, respectively. We observed that the nontransmitted parental PMMs have a distinct AF distribution, which is bimodal, and significantly different from both transmitted parental PMM and child PMM distributions (nontransmitted parental v. transmitted, p = 7.07 × 10^−14^, nontransmitted parental v. children, p = 2.99 × 10^−14^, two-sided WRST, FDR <0.05). Our analysis of transmission filtering suggests that a major fraction of calls from the rightmost portion of the distribution are false mosaic calls that would have been filtered given the availability of grandparental data (Figure S21). Interestingly, however, the parental transmitted PMM distribution closely resembles the rightmost mode of the nontransmitted distribution, suggesting that this subset is still representative of likely early embryonic events, a fraction of which are also found in the germ cells. We further examined the AFs of the parental PMMs taking into account the confidence intervals of the AFs, similar to how we empirically separated germline and mosaic calls (Figure S23). Although we found some transmitted variants within the low AF range, the vast majority had AF CIs in excess of 10% (92/94 [98%]), suggesting early embryonic origin for PMMs within this AF range and consequently the largest risk for transmission.

**Figure 4.**
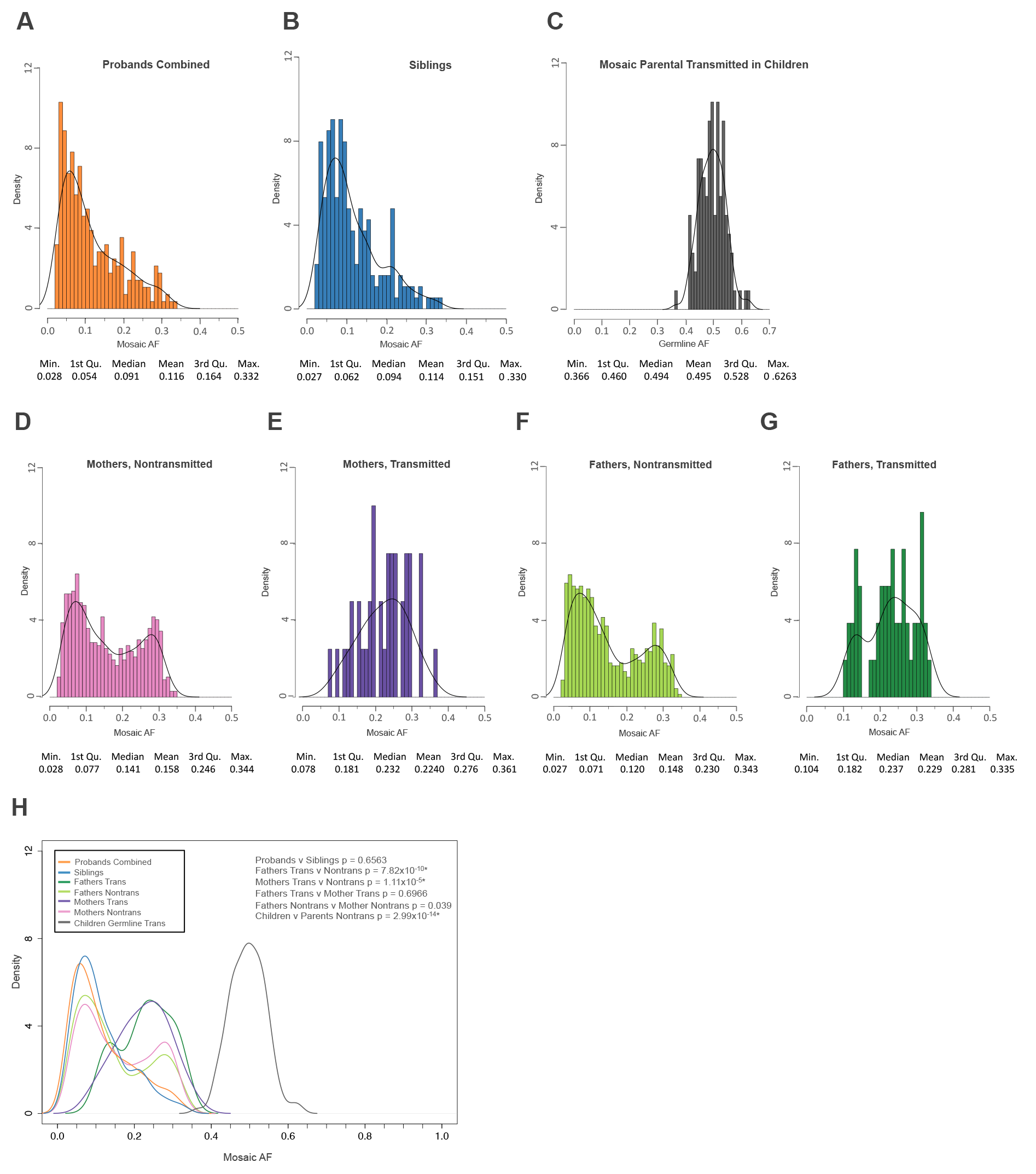
Mosaic Variant Allele Fraction Distributions. For all plots, all PMMs from the 5%-45× high confidence call set were used. (A) Distribution of allele fractions for variants in probands combined (quad + trio families).
(B) Distribution of allele fractions for variants in siblings.
(C) Distribution of allele fractions for germline variants in children that were transmitted from mosaic parents.
(D) Distribution of allele fractions for variants in mothers that were not transmitted to children.
(E) Distribution of allele fractions for variants in mothers that were transmitted to children.
(F) Distribution of allele fractions for variants in fathers that were not transmitted to children.
(G) Distribution of allele fractions for variants in fathers that were transmitted to children.
(H) Combined data plotted as kernel density curves. Parental transmitted are significantly shifted towards a higher allele fraction than nontransmitted or child mosaic variants. Children have a significantly different distribution than parental nontransmitted. Significance determined using a two-sided Wilcoxon rank sum test. *FDR < 0.05 using the Benjamini-Yekutieli approach.

We also examined the mutational spectra of GDMs and PMMs. As has been previously demonstrated,^15^ we found the majority of mutations are transitions over transversions for both GDMs and PMMs (parents and children), with the most common mutation being C>T transitions at CpG dinucleotides (Figure S24A). The relative frequency of mutations within trinucleotides showed strongest correlation with previously described cancer signature 1,^40^ for both GDMs and PMMs (Figures S24B-C). Signature 1, which is characterized by spontaneous deamination of 5-methylcytosine, is indicative of endogenous mutational processes and associated with all cancer types.^40^ Unlike previous results,^15^ our second strongest correlated cancer signature was 6, which is associated with defective DNA mismatch repair.^40^

### Potential Impact of Synonymous PMMs on Splicing

We hypothesized that if the observed burden of synonymous PMMs contributed to ASD risk, one possible mechanism would be by disrupting splicing within the associated exon. If this was the case, we expected proband synonymous PMMs to be preferentially localized near existing canonical splicing sites, i.e. the starts or ends of exons.^56; 57^ Therefore, we calculated the absolute minimum distances of all synonymous PMMs and GDMs to their closest splicing site (Figure 5). We found the proband synonymous PMM distribution to be shifted towards splicing sites compared to both sibling and parental synonymous PMM distributions (p = 0.017 and p=0.008 respectively, two-sided WRST, FDR < 0.05), while the sibling distribution was similar to the parental (p = 0.61, two-sided WRST). We observed a similar shift towards splice sites for GDMs in probands as compared to siblings (p = 0.005, two-sided WRST, FDR < 0.05).

**Figure 5.**
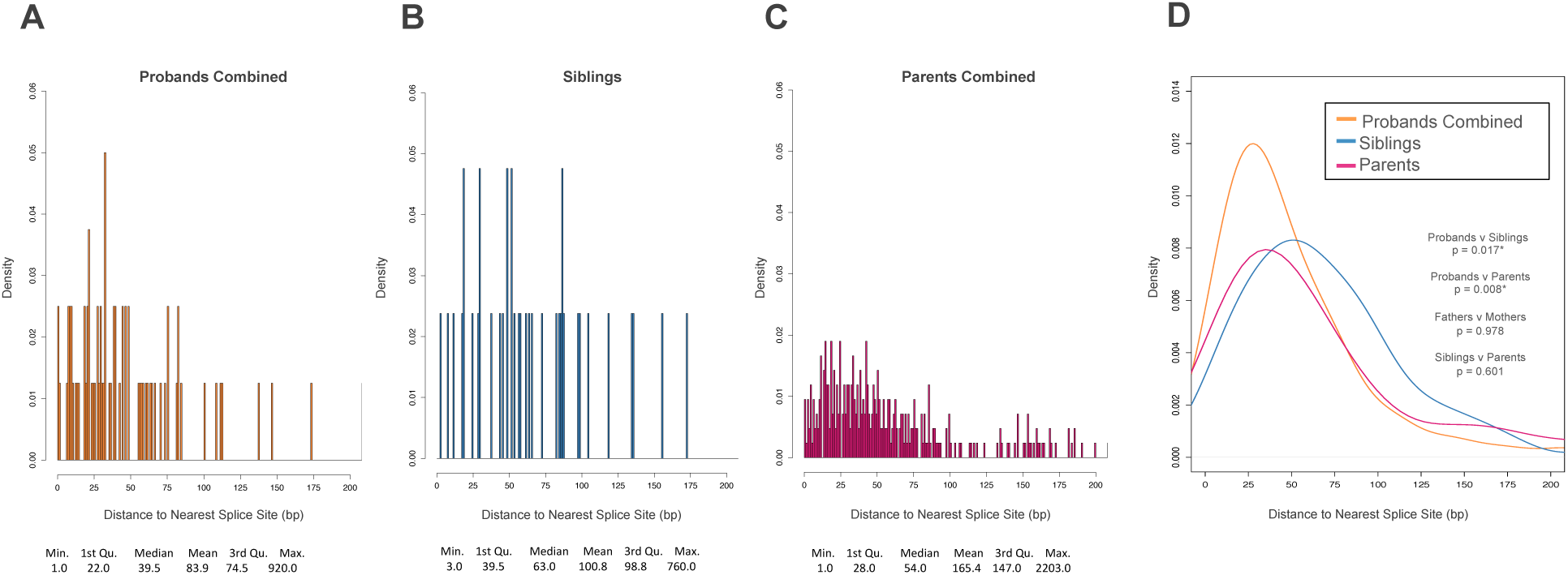
Distance to Nearest Splice Site for Synonymous PMMs. For all plots, all synonymous PMMs from the 5%-45x high confidence call set were used. Splice site distances were calculated as absolute minimum distance to nearest canonical splice site. (A) Distribution of distance to nearest splice site in probands combined (quad + trio families).
(B) Distribution of distance to nearest splice site in siblings.
(C) Distribution of distance to nearest splice site in combined parents (quad + trio families).
(D) Combined data plotted as kernel density curves. Proband distribution is significantly shifted towards the canonical splice sites compared to both parents and siblings. Significance was determined using a two-sided Wilcoxon rank sum test. *FDR < 0.05 using the Benjamini-Yekutieli approach.

We further evaluated potential effects of synonymous mutations on splicing computationally using HSF.^41^ HSF utilizes a collection of splicing prediction approaches that assess disruptions of known splicing factor binding motifs through: position weight matrix analysis, comparing presence of the mutant or wild type sequence in multiple putative splice regulation motif collections, and performing maximum entropy modeling. HSF reported significantly more instances of putative splice altering mutations for proband synonymous PMMs (70/78) when compared to siblings (25/41) (p = 0.0005, odds ratio, 5.506, 95% CI 1.946-16.836, two-sided Fisher’s exact) (Table S6). Synonymous GDMs showed no enrichment (proband-188/235 v. sibling-137/177, p = 0.544, odds ratio, 1.168, 95% CI 0.726-1.879 two-sided Fisher’s exact). When restricting to synonymous PMMs that occur within 50 bp of the start or end of an exon, where splicing regulatory elements are enriched^58^, we observed a stronger enrichment (proband-45/53 v. sibling-5/12, p = 0.00378, odds ratio, 7.53, 95% CI 1.618-38.861, two-sided Fisher’s exact). We did not observe a similar enrichment for proband synonymous GDMs near splice junctions. To assess the robustness of HSF findings, given the high call rate of splice altering variants, we removed the two most frequently called matrices and reclassified variants. We still observed an enrichment of proband synonymous PMMs predicted to alter splicing (all variants: proband-53/79, sibling-18/41, p = 0.019, odds ratio, 2.60, 95% CI 1.20-5.66; within 50 bp: probands-34/50, sibling-5/15, p = 0.033, odds ratio, 4.25, 95% CI 1.24-14.5, two-sided Fisher’s exact).

To independently assess splice altering variant enrichment, we applied a recently reported machine-learning based approach, SPANR.^42^ SPANR requires a variant to be within 100 bp from an exon start or end site and be located within an exon flanked by an exon on either side. This limited our SPANR analysis to 68 proband and 29 sibling PMMs. SPANR reported a significant enrichment of splice altering synonymous PMMs in probands (proband-15/68, v. sibling 1/29, p = 0.03, odds ratio, 7.81, 95% CI 1.09-344.8, two-sided Fisher’s exact). Similarly, proband PMMs remained enriched for splice altering variants (though not significantly) when restricting to mutations within 50bp of a canonical splice site (proband-14/46, sibling-1/13, p = 0.15, odds ratio 5.13, CI 95% 0.64-239.9, two-sided Fisher’s exact).

### Geneset Enrichment

We applied a similar approach as Iossifov and colleagues to look for enrichments of PMMs within different gene sets using our high confidence dataset.^11^ Using expected values generated from joint coverage for the cohort, we examined whether our PMMs/GDMs showed more or less mutations than expected independently for probands and siblings. As expected, our GDM dataset showed similar enrichments or lack thereof to previous reports (Table 3). In probands, we found enrichment (1.8-fold) for missense PMMs intersecting chromatin modifiers (p = 0.043, two-sided binomial) and depletion of missense PMMs in embryonically expressed genes (p = 0.024, two-sided binomial). Interestingly, missense GDMs showed no evidence of enrichment or depletion for these gene sets, while LGD GDMs have previously been shown to be enriched.^11^

**Table 3.** Enrichment of Missense Germline and Postzygotic Mutations in Gene Sets

|  |  |  | Mis GDM (Pro) |  |  | Mis GDM (Sib) |  |  | Mis PMM (Pro) |  |  | Mis PMM (Sib) |  |  |
| --- | --- | --- | --- | --- | --- | --- | --- | --- | --- | --- | --- | --- | --- | --- |
| Total no. of genes= |  |  | 701 |  |  | 426 |  |  | 177 |  |  | 129 |  |  |
| Set | *p* | Genes in set<sup>a</sup> | Obs | Exp | *P* | Obs | Exp | *P* | Obs | Exp | *P* | Obs | Exp | *P* |
| **Chromatin** | 0.0372 | 388 | 32 | 26.1 | 0.230 | 20 | 15.8 | 0.303 | 12 | 6.6 | 0.043 | 2 | 4.8 | 0.247 |
| **Embryonic** | 0.1433 | 1,797 | 114 | 100.5 | 0.178 | 60 | 61.1 | 0.835 | 16 | 25.4 | 0.024 | 25 | 18.5 | 0.103 |
| **Essential** | 0.1967 | 2,402 | 160 | 137.8 | 0.036 | 83 | 83.7 | 0.903 | 41 | 34.8 | 0.256 | 24 | 25.4 | 0.825 |
| **PSD** | 0.0701 | 879 | 58 | 49.1 | 0.183 | 35 | 29.9 | 0.346 | 17 | 12.4 | 0.183 | 14 | 9.0 | 0.167 |
| **FMRP** | 0.1005 | 775 | 100 | 70.3 | 4x10<sup>-4</sup> | 57 | 42.7 | 0.036 | 20 | 17.8 | 0.53 | 13 | 12.9 | 1.000 |

45x joint coverage, 5% AF call set. Variants in sex chromosomes excluded. Expected (Exp) and *P* values obtained from two-sided binomial test, based on gene length model (*p*). Abbreviations: Mis GDM-missense germline *de novo* mutation, Mis PMM-missense postzygotic mutation, Pro-proband, Sib-sibling, PSD-post synaptic density associated genes, FMRP-fragile X mental retardation protein-associated genes. <sup>a</sup>Total number of genes differs from full lists as only genes that we were able to map to our gene symbol annotations were used and genes on sex chromosomes were excluded.

Recently, several groups have taken different approaches to generate genomewide ASD candidate risk gene rankings and predict novel gene targets.^36; 43^ These approaches have largely been validated on LGD GDMs. We explored whether our high confidence PMM calls showed any shift in ASD candidate gene rankings for probands compared with their unaffected siblings (Table S16). We evaluated rankings based on gene mutation intolerance (LGD rank, LGD-RVIS average rank)^36^ or based on a human brain-specific gene functional interaction network (ASD association).^43^ At the population level, we found only non-significant increases in LGD-RVIS rankings for proband synonymous and essential missense PMMs in the subcohort of families without any proband NS GDMs (p = 0.029 and p = 0.073, one-sided WRST). We also observed no significant shifts in rankings for missense GDMs.

### Targets of Proband Mosaic Mutations

To determine if germline and mosaic mutations in probands share common targets, we intersected missense PMMs from the high confidence call set and the burden subset (15%-45×) with the re-classified published GDM calls. We observed no enrichment of proband missense PMMs with genes that are targets of sibling GDMs of any type. However, we did find an apparent enrichment of genes that are targets of proband missense GDMs within proband missense PMMs from the burden call set (proband: 25/100; sibling: 9/69, p = 0.042, OR, 2.222, 95% CI 0.904-5.582, one-sided Fisher’s exact), suggesting some common ASD risk targets for mosaic and germline mutations.

We next examined the phenotypic features of probands with PMMs from our high confidence and re-classified published call sets. We focused on probands with NS mutations that intersect genes with previously published NS GDMs, as these genes have higher prior probabilities for being involved in ASD risk.^11^ We compared a range of clinical indices among 12 subjects with LGD PMMs (mean age: 9.9 years, standard deviation [SD] 3.8; 83.3% male) and 45 subjects with missense PMMs (mean age: 9.5 years, SD 3.9; 81.8% male) (Supplemental Material and Methods and Supplemental Note: Case Reports). We found no between-group differences with respect to developmental features, autistic symptomatology, or non-autistic behavioral and emotional problems. Level of functioning, however, appeared to be slightly more impaired in the LGD PMM group. This was evidenced with Vineland standardized scores that were significantly lower in the LGD versus the missense group for the Adaptive Behavior Composite (66.4 v. 73.8, p = 0.037, t-test) and the Social Interpersonal domain score (64.5 v. 72.3, p = 0.041, t-test) whereas differences in standard scores for the Communication (69.5 v.77.0, p = 0.057, t-test) and Daily Living Skills (69.8 v. 77.6, p = 0.076, t-test) domains showed non-significant trends. Intellectual functioning showed differences of the same magnitude and direction. Full scale IQ showed a mean difference of 18.1 points (66.2 v. 84.3, p = 0.060, t-test) that was not significant largely due to the large standard deviations of IQ scores in both groups (26.8 and 29.5, respectively). Differences for nonverbal IQ (71.8 v. 85.4, p = 0.142, t-test) and verbal IQ (66.7 v. 83.8, p = 0.099, t-test) were of the same magnitude but were not statistically significant.

In addition, we intersected all predicted NS PMMs (our high confidence call set plus reclassified published [unique CDS]) with 171 genes that have reached high confidence levels for their risk contribution for ASD and developmental disorders from large WES and targeted resequencing cohorts.^11; 35; 45; 46^ In probands, 15/496 PMMs intersect (9 missense, 6 LGD) while only 4/354 PMMs intersect in siblings (3 missense and 1 LGD). The novel, i.e. not published in the GDM call set (Table S14),^11; 12^ proband events included missense PMMs in: *CHD2* (MIM: 602119, p.Glu91Gly), *CTNNB1* (MIM: 116806, p.Arg376His), *RELN* (MIM: 600514, p.Gly3239Arg), *KIF1A* (MIM: 601255, p.Ala219Thr), and *KMT2C* (MIM: 606833, p.Arg4806Gly) (Table 4). We also identified a novel missense mutation in *SCN2A* (MIM: 182390, p.Ser1124Cys) that was transmitted to the proband from the mother. Our SNV PMM pipeline reidentified published *de novo* calls that we re-classified as likely mosaic events, including: *KANSL1* (MIM: 612452, p.Gln243His), *KAT2B* (MIM: 602303, splicing), *INTS6* (MIM: 604331, p.Arg596Ter), *SYNGAP1* (MIM: 612621, p.Arg1019Cys), and *TBL1XR1* (MIM: 608628, p.Leu282Pro) (Table 4). Only the *KANSL1* and *INTS6* PMMs met the high confidence 45× joint coverage criteria. Mosaic re-classified indels included: *DIP2A* (MIM: 607711, p.Leu552ValfsTer34), *GIGYF1* (MIM: 612064, p.Thr381ArgfsTer13), *SRCAP* (MIM: 611421, p.Asp2137GlufsTer25), and *ZC3H4* (MIM: NA, p.Lys449ThrfsTer12) (Table 4). With the exception of probands with the *CHD2* and *DIP2A* PMMs, none of these other probands have NS GDMs in other strong risk genes.

**Table 4.**
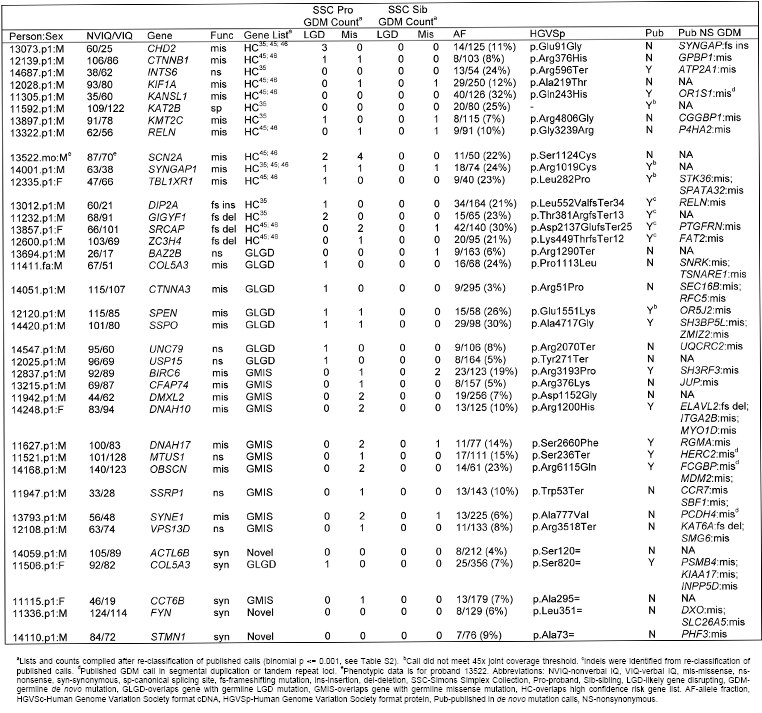
Highlighted Mosaic Mutations In Candidate ASD Risk Genes

Among the remaining NS PMMs, we found seven mutations in genes overlapping proband LGD GDMs (sibling NS GDM count <= 1) (Table 4). Of particular interest are novel nonsense PMMs in *BAZ2B* (MIM: 605633, p.Arg1290Ter), *UNC79* (MIM: 616884, p.Arg2070Ter) and *USP15* (MIM: 604731, p.Tyr217Ter). *BAZ2B* is part of the bromodomain gene family involved in chromatin remodeling.^59^ *UNC79* works in concert with *UNC80* to regulate the excitability of hippocampal neurons through activation of sodium channel NALCN.^60^ *USP15* is a deubiquitinase that plays many roles across the cell including modulating immune response through TGF-β and NF-KB pathways.^61^

Ten of the remaining NS PMMs intersect gene targets of missense GDMs (sibling NS GDM count <= 1) (Table 4). Of note are novel nonsense PMMs in the chromatin remodeling factor *SSRP1* (MIM: 604328, p.Trp53Ter) and the membrane trafficking protein *VSP13D* (MIM: 608877, p.Arg3518Ter). Novel missense PMMs included: *DMXL2* (MIM: 612186, p.Asp1152Gly), *SYNE1* (MIM: 608441, p.Ala777Val), and *CFAP74* (p.Arg376Lys).

Among the synonymous PMMs, we identified four novel candidate genes based on known roles in neurodevelopment, predicted creation of a new exonic silencing site, and no other NS GDM events in ASD risk genes in the proband: *ACTL6B* (MIM: 612458, p.Ser120=), *CCT6B* (MIM:610730, p.Ala295=), *FYN* (MIM: 137025, p.Leu351=), and *STMN1* (MIM: 151442, p.Ala73=). Notably, *ACTL6B* is a neuron-specific component of the SWI/SNF chromatin remodeling complex.^62^ We also highlight a synonymous PMM in *COL5A3* (p.Ser820=) as: it has a high likelihood of impacting splicing by altering the wild type 3’ exonic donor site, a missense PMM (p.Pro1113Leu) and a LGD GDM are present at this locus, and we found no other NS GDMs associated with ASD risk in the proband. Taken together, these new mosaic calls provide additional support for high confidence ASD risk genes and highlight novel candidates as potential contributors to ASD risk.

## Discussion

The aim of our study was to systematically evaluate exonic PMMs in a large-family based SSC cohort and their potential role in ASD. Historically, PMMs, much like GDMs, have been intractable to systematically study genomewide. However, NGS technologies have now made this class of genomic variation accessible, genomewide, at single base resolution. A number of recent reports have demonstrated that PMMs are relatively common in both healthy and neurodevelopmental disorder cohorts, including intellectual disability, ASD, or general developmental delays. ^2; 15; 27; 63; 64^ However, how frequent and widespread these events might be in early and/or late development and how much risk they contribute to complex disorders has yet to be fully elucidated.

We began by analyzing the underlying variant support data for previously reported *de novo* mutations from the SSC WES data. These calls were generated using a variety of calling approaches designed to identify germline variants. We found evidence for 11% of SNVs and 26% of indels having AFs consistent with a PMM arising in the child. This is in excess of our original observation of 3.5% (9/260) of mutation events consistent with child PMMs, using only 209 families.^2^ A similar analysis of *de novo* mutations identified from whole-genome sequencing of simplex ID trios validated 6.5% (7/107) as PMMs.^27^

As the previously published SSC *de novo* calls were all generated with germline variant callers, we reasoned that re-analyzing the WES data systematically with approaches tuned to detect PMMs would reveal novel mutations, especially those with lower AFs (<20%). Therefore, we sought to develop generalizable methods to identify PMMs and GDMs that can be applied to NGS data, without matched normal data but in the context of nuclear families. We focused our analysis on SNVs because of their higher predicted frequency and more uniform AF distribution. We developed our method by integrating calls using three complementary approaches and performing extensive validations using smMIPs (Figure 1E). Using these data, we developed a logistic regression model and additional heuristics that clearly separate false calls, PMMs, and GDMs. Given that the depth of sequence directly affects the observable minimum mutation AF, we used varying AF-COV thresholds (e.g. 15%-45×, 5%-130×) to evaluate mosaic mutation burden. Surprisingly, in the full cohort, we found the strongest signal for PMM burden with synonymous SNVs (Figure 2C). The distribution of proband PMMs showed a significant shift in distance to nearest splice site (Figure 5D). Moreover, proband synonymous PMMs showed enrichments for splice altering predictions using two independent approaches.

It has recently been shown that in some cancers synonymous mutations may have a modest enrichment in oncogenes.^56^ Within 16 oncogenes, the signal was specific to the mutations within 30 base pairs (‘near-splice’) of the exon boundary and showed gains of exonic splicing enhancer (ESE) motifs and loss of exonic splicing silencer (ESS) motif sequences. Conducting an analysis of the intersection of ASD and schizophrenia WES GDMs and regulatory elements, Takata and colleagues recently reported an enrichment of near-splice synonymous GDMs in ASD probands (odds ratio ∼2) and, to a lesser extent schizophrenia probands, relative to controls.^57^ Stronger signal in their initial ASD cohort was seen for sites predicted to cause ESE/ESS changes, but reduced in a replication dataset (odds ratios 2.52 and 1.55 respectively). In their analysis they compared the fraction of near-splice or those also disrupting ESE/ESS sites mutations in cases versus controls (Fisher’s exact test), which does not take into account coverage differences across individuals/cohorts. We repeated our analysis of the distance to splice site distributions for the high confidence 45x-joint coverage SSC synonymous GDMs, finding them to be significantly closer to splice sites in probands as compared to siblings (p = 0.005), similar to the PMM calls. However, we observed no corresponding enrichment of splice altering variant predictions. Taken together, these data are consistent with a possible role of synonymous postzygotic mutations that functionally disrupt splicing regulation in ASD.

While computational splice regulation predictions can provide useful information at the population level, we advise interpreting the effect of individual variants with caution given the uncertainty of splice regulatory mechanisms, cell-type specific splicing patterns, limited training sets, and high reported false positive rates. For example, HSF has a reported false positive rate (43%).^41^ This is due in part to the wide breadth of splicing signals it attempts to capture. Additional functional validation of these mutations using *in vitro* approaches, e.g. minigene assays, or *in vivo* approaches, e.g. genome editing of cell lines, is warranted.

From the synonymous PMMs predicted to impact splicing, we identified a number of genes that have roles in neurodevelopment and are associated with other ASD risk genes. Additionally, these events are within probands who either have no other GDMs or whose other GDMs would not likely be associated with ASD risk. In particular, we highlight genes *ACTL6B*, a member of the chromatin remodeler complex SWI/SNF;^62^ *CCT6B*, a postsynaptic density gene recently implicated in recessive intellectual disability;^65^ *FYN*, which encodes a non-receptor tyrosine kinase that is involved in axon outgrowth;^66^ and *STMN1*, which encodes a microtubule destabilizing protein that is involved in the regulation of axon outgrowth.^67^ Also notable is *COL5A3*, which encodes a scaffolding protein that is directly regulated by ASD and Pitt-Hopkins (MIM: 610954) associated gene *TCF4* (MIM: 602272).^68^ Individuals with duplications that span *COL5A3* have phenotypic characteristics similar to those of TCF4-related syndromes including seizures, facial dysmorphia, and developmental delay.^68^

We did not observe evidence of missense PMM burden in the full cohort of ASD probands. This is perhaps not surprising given the strong contribution of GDMs to ASD in the SSC and that most *de novo* events will be missense changes by chance, i.e. form most of the background non-disorder related mutations. Our sample size is too small given their rate of mutations to fully evaluate nonsense/splice PMMs as a separate class. Based on the differential between probands and siblings, it has been reported that LGD GDMs have a 40% likelihood of being contributing to ASD (90% of loci with recurrent LGD), while the likelihood for missense variants is ∼35%.^11^ We reasoned that restricting our analysis to families without proband germline mutations would increase our power to detect any effect of missense PMMs, even though we would be removing a significant fraction of families with germline events unrelated to ASD. Indeed, if we subdivide the SSC cohort into families that have or do not have a proband LGD GDM/de *novo* CNVs, or, alternatively, any NS germline mutation, we observed a difference emerging. This difference is strongest in the subset of genes predicted to be essential/intolerant to mutation (Figure 3B and 3C). Similarly, we also saw a further increase in synonymous PMM burden in the subcohort without any reported NS GDMs (Figure 2).

Freed and Pevsner recently reported on PMM burden in probands and siblings in the SSC.^63^ While our two studies used the same SSC datasets, we each used different computational and validation approaches. Our 45× high confidence SNV call set contains 470 PMMs in children, 398 that are unique to our study. Their 20x high confidence SNV call set contained 196 PMMs, 124 that are unique (65 calls <40× coverage). Most importantly, they restricted their burden analysis to their PMM calls that: overlapped the previously published *de novo* datasets, met 40x joint-coverage, and also included indel calls. Unlike our study, they did not restrict their analysis to different minimum AF-COV thresholds. They report the burden of all classes of variants combined (synonymous, missense, and LGD) as significant and estimate that 5.1% of probands have PMMs related to ASD risk. Moreover, they found nominal contributions across all classes of mutations. Comparing our 15%-45× PMM analysis to their data, we observed similar differences in the synonymous rates, but not their observed missense differences in the full cohort. These differences are likely driven by our different computational approaches and our use of a larger number of PMM calls unique to our pipeline. Specifically regarding the burden analyses, 175/231 of our SNV PMM calls are unique to our study. Of the 131 re-classified published SNV calls included in the Freed and Pevsner burden analysis, 75 were absent from our call set because they failed best practice filters, did not meet minimum joint coverage (40× v. 45×), or were present in excluded families. With our approach, we estimate that PMMs as a group contribute to 4-8% of simplex ASD, with an ∼2% contribution from synonymous mutations. Combined our two analyses suggest that exonic PMMs as a whole are likely contributing to ASD risk in the SSC at rates similar to other classes of *de novo* mutations.^11; 35^

We also tested whether PMMs and GDMs shared common gene targets. We found proband missense PMMs were more likely than sibling missense PMMs to intersect with proband GDMs (odds ratio ∼2). We also found that a number of our novel nonsense PMMs in probands overlapped genes with GDMs including: *BAZ2B, SSRP1, UNC79, USP15*, and *VPS13D* (Table 4). Consistent with our observation of enrichment of chromatin modifiers in proband missense PMMs, we found many of our PMMs overlapping genes with NS GDMs are also involved in chromatin regulation: e.g., *BAZ2B, CHD2, COL5A3, KAT2B, KMT2C*, and *SSRP1*. Recent studies have found that ASD risk genes are highly co-expressed during the mid-fetal period of cortical development.^69; 70^ Several PMMs intersect genes that occupy the same co-expression modules, which are significantly enriched for ASD risk genes. For example, *BIRC6* (MIM: 605638), *DMXL2, OBSCN* (MIM: 608616), *SPEN* (MIM: 613484), *SRCAP, SSRP1, UNC79*, and *ZC3H4*, all occupy modules 2 and 3, which peak between post conception weeks 10-22 and are enriched for chromatin modifiers/transcriptional regulators.^69^ *COL5A3, KIF1A, SCN2A*, and *SYNE1* are found in modules 13/16/17, which are turned on later in development, after post conception weeks 10, and are enriched for synaptic genes.^69^

Moreover, we found missense PMMs in some of the highest confidence ASD risk genes identified in the SSC or other combined studies, for example: *CHD2, CTNNB1, KMT2C, SCN2A* and *SYNGAP1* (Table 4).^33; 35; 36; 71^ Interestingly, small *de novo* deletions targeting *CHD2, SYNGAP1, CTNNB1, and KMT2C* have been reported in the SSC as well,^35^ demonstrating that new mutations of multiple types and origins at these sites contribute to ASD risk. Taken together our data argues that proband PMMs and GDMs target many common risk genes. Finally, mutations in some of these genes are not restricted to ASD as these genes have also been found to be disrupted in cohorts primarily defined on diagnoses of: epileptic encephalopathy, ID, and congenital heart defects with additional features.^72–75^ Understanding how mutations impact these important genes that blur our diagnostic constructs will be an important area of future research. These and other data suggest the creation of more broadly defined cohorts and better integration of genetic studies of developmental disorders are warranted.

Unique to this study, we also performed our PMM analyses in the parental data. We identified both nontransmitted and transmitted PMMs. Transmitted PMMs are obligated to be present in both the soma and the germline. Given the low number of offspring of each parent, we cannot rule out the possibility that a fraction of the nontransmitted parental events are also present in the parental germ cells. Our observed postzygotic mutation rate is much higher in the SSC parents compared to the SSC children. Moreover, the nontransmitted PMM AFs have a bimodal distribution that is distinct from both the child PMMs and parental transmitted PMMs. There are several potential explanations for the increased rate of mutation and AF differences. As parents in this cohort were several decades older at time of DNA collection, this increase could be explained by the accumulation of PMMs in the blood, some of which might drift to or be selected for higher AF. We found very little evidence for enrichment of PMMs in genes related to blood ACEs, except *DMNT3A*. The number of parents with PMMs in ACE-related genes is <1%, which is consistent with estimates that ACE associated mutations occur in less than 1% of individuals under 50 and do not begin to rise until after 65.^53–55^

We did not evaluate the grandparental generation. For the children, we were able to eliminate likely inherited calls that showed consistent bias in AF generation to generation (about two-thirds of filtered calls) as well as some additional calls that exhibited skewing likely due to random chance (Figure S6). Our analysis on a subset of the cohort suggests that ∼40% of the excess in nontransmitted parental PMM calls could be explained by incomplete filtering of these recurrently biased and randomly skewed sites, while the remained are likely true events (Figure S21). Based on the children, recurrently biased sites are likely to have higher AFs (> 20%). PMMs with AF that fall in this upper range that are not clearly transmitted should be interpreted with caution. Importantly, Xie and colleagues report this same bimodal distribution in a case-control study of ACE, which did not benefit from transmission based filtering.^54^

Rahbari and colleagues recently performed whole-genome sequencing on moderately sized pedigrees followed by the identification and characterization of *de novo* mutations in multiple children, spanning approximately a decade.^15^ We similarly observed that the cancer mutation signature 1^40^ is most strongly correlated with *de novo* mutations (GDMs and PMMs). This signature is associated with endogenous mutational processes. Our second strongest signature was 6, which is associated with defective DNA mismatch repair. Rahbari and colleagues in contrast identified signature 5, which has no known etiology but is common to all cancer types. This finding could reflect differences between the content of WES versus whole-genomic datasets. In validating their *de novo* calls using target capture and deep sequencing, they identified a number of mutations that were at low levels in the parental blood derived DNA. Importantly in contrast to our study, PMMs were not directly identified in the parents and calls with greater than 5% of reads showing the alternative allele in a parent were excluded from the *de novo* call set. Nevertheless, they found that 4.2% of apparent germline mutations are present in the blood of parents at >1% AF. However, the rate we observed in our high confidence smMIP validation data, of similar calls (without parental WES signal), is 0.6% (1 out of 164). In our 45× WES dataset, we found 0.66% of GDMs in children are also obligate gonadal mosaic. Overall, our data support at least 7-11% (depending on the AF) of parental PMM events are also present in the parental germ cells and can be transmitted to the next-generation. Together these two sets of parental postzygotic mutations account for 6.8% of all germline mutations present in the children from our high confidence call set (Table S5). Importantly, many of these events would be missed by *de novo* calling pipelines that eliminate any sites with variant reads present in a parent. This rate is higher than what has been recently reported for *de novo* CNVs (4%).^23^ These findings have important implications for recurrence risk and clinical testing, which are still not widely appreciated.^14; 15; 23; 76; 77^ While the recurrence risk for *de novo* mutations is generally thought to be low (∼1%), finding the presence of a mutation, even at low levels, in a parent dramatically increases this risk to a previously estimated >5%.^15; 76; 77^ The risk may be dramatically higher for specific mutations, depending on their embryonic timing and distribution within the germ cells.

We were limited by the availability of DNA from a single peripheral blood source and WES data that is non-uniform. Future studies in this area would greatly benefit from deep uniform whole-genome sequencing, access to multiple peripheral and other tissue types of different embryonic origin, and improved indel variant calling approaches. This could include brain tissue in cases of surgical resection to control intractable epilepsy. Moreover, we strongly suggest that new efforts to establish autism brain banks obtain peripheral DNA samples from the donor and their parents. These DNA would greatly aid in the classification of variant types,i.e. PMMs, GDMs, or inherited variants, identified in bulk brain and single-cell sequencing studies as well as help determine their likely embryonic timing.

In summary, our data support the conclusion that exonic postzygotic mosaicism contributes to the overall genetic architecture of ASD, in potentially 4-8% of all ASD simplex cases, and that future studies of mosaicism in ASD and related-disorders are warranted. We present a general approach for identifying PMMs that overcomes many of the inherent detection and validation challenges for these events in family-based and unmatched samples. The methods developed will allow continued discovery of PMMs in future datasets, including unsolved genetic disorders, and our findings have potential translational implications for clinical detection, case management, interventions, and genetic counseling.

## Supplemental Data

Supplemental Material and Methods

Supplemental Note: Model Development

Supplemental Note: Case Reports

Supplemental References Figures S1-S24

Tables S1-S17

## Acknowledgments

This work was supported by a grant from the Simons Foundation (SFARI 305927, B.J.O) and the Agence Nationale de la Recherche (ANR-13-PDOC-0029, Y.D. and J.-B.R.). B.J.O. is currently a Klingenstein-Simons Fellow in Neurosciences, Alfred P. Sloan Foundation Fellow in Neuroscience, and is supported by the NARSAD Young Investigator Award from the Brain and Behavior Research Foundation. We are grateful for the use of the Exacloud high performance computing environment developed jointly by OHSU and Intel and the technical support of the OHSU Advanced Computing Center. We would like to thank S.J. Webb, A.C. Adey, K.M. Wright, I. Iossifov, S. Bedrick, J. Burchard, A. Presmanes Hill for helpful discussions regarding the manuscript. We also thank I. Fisk, N. Volfovsky, N. Krumm, and T.N. Turner for their assistance accessing the WES datasets. We are grateful to all of the families at the participating Simons Simplex Collections (SSC) sites, as well as the principal investigators (A. Beaudet, R. Bernier, J. Constantino, E. Cook, E. Fombonne, D. Geschwind, R., Goin-Kochel, E. Hanson, D. Grice, A. Klin, D. Ledbetter, C. Lord, C. Martin, D. Martin, R. Maxim, J. Miles, O. Ousley, K. Pelphrey, B. Peterson, J. Piggot, C. Saulnier, M. State, W. Stone, J. Sutcliffe, C.Walsh, Z. Warren, E. Wijsman). Approved researchers can obtain the SSC population dataset described in this study by applying at https://base.sfari.org.

## Web Resources

The URLs for data presented herein are as follows:

UCSC Genome Browser, http://genome.ucsc.edu/

OMIM, http://www.omim.org

GenBank, https://www.ncbi.nlm.nih.gov/genbank/

SFARI base, https://base.sfari.org.

Simulated NGS Data, http://www.ebi.ac.uk/goldman-srv/simNGS/

Variant Effect Predictor, http://uswest.ensembl.org/Tools/VEP

ANNOVAR, http://annovar.openbioinformatics.org/

GenPhenF, https://iossifovlab.com/gpf

Mutational Signatures Data, http://cancer.sanger.ac.uk/cancergenome/assets/signatures_probabilities.txt

Human Splice Finder, http://www.umd.be/HSF3/SPANR, http://tools.genes.toronto.edu/

